# CCR8⁺ regulatory T cell depletion overcomes immune checkpoint blockade resistance in BRCA2-deficient prostate cancer

**DOI:** 10.64898/2026.11.11.765808

**Authors:** Kimberly A. Rickman, Irena Q. Sun, Ramya Parameswaran, Xiaolin Zhu, Elise Y. Cai, Valentina Rossi, Nicole Ziper, Hanbing Song, Christina Thai, Yingyue Li, Morgan E. Diolaiti, Franklin W. Huang, Harish N. Vasudevan, Alan Ashworth

## Abstract

*BRCA2*-mutated prostate cancer is an aggressive, genomically unstable subtype, yet how this genomic instability influences tumor immunity remains undefined. We developed an isogenic, immunocompetent model of *Brca2*-deficient prostate cancer in which BRCA2 loss activated the cGAS/STING pathway and a type I interferon response across multiple cellular compartments, paradoxically leading to an overall immunosuppressed tumor microenvironment. Within this microenvironment, tumor-associated macrophages were interferon-responsive and expressed the CCR8 ligand *Ccl8*, and there was enrichment of immunosuppressive CCR8⁺ regulatory T cells (Treg). Anti-PD-1 monotherapy was ineffective and instead reinforced this immunosuppressive circuit, leading to expansion of CCR8⁺ Tregs. However, depletion of CCR8⁺ regulatory T cells overcame resistance to checkpoint blockade. Analysis of human *BRCA2*-mutated prostate cancer showed elevated *CCR8* expression, supporting the translational relevance of the CCR8⁺ Treg axis as a potential therapeutic target in a disease largely refractory to immunotherapy.

## Introduction

Alterations of DNA damage repair (DDR) genes define a clinically high-risk subset of prostate cancers, accounting for approximately 20% of advanced disease, and the most commonly mutated gene is *BRCA2*, followed by *ATM*, *CDK12*, and *CHEK2.*^1–5^ *BRCA2*-mutated tumors have an aggressive natural history characterized by higher grade, more advanced disease at diagnosis, and higher recurrence rates after definitive therapy for localized disease.^6–9^

The primary function of BRCA2 is in homologous recombination (HR)-mediated repair of DNA double-strand breaks (DSBs).^10,11^ BRCA2 loss results in genomic instability and defective repair of replication-associated DNA damage.^10,12^ These DNA repair pathway defects underlie sensitivity to PARP inhibitors (PARPi), which trap PARP1 on DNA and impair the repair of single-strand breaks, generating replication-associated lesions that require HR for resolution.^13^ PARPi were originally approved for metastatic castration-resistant prostate cancer (mCRPC) with DNA repair defects.^14,15^ More recently, PARPi in combination with an androgen receptor pathway inhibitor (ARPI) received FDA approval for *BRCA2*-mutated metastatic castration-sensitive prostate cancer (mCSPC).^16,17^ Despite substantial clinical benefit, resistance remains inevitable, underscoring the need for improved treatments for high-risk DDR-altered tumors, including approaches that exploit tumor-immune interactions rather than DNA repair vulnerabilities alone.

Immune-checkpoint blockade has shown only limited efficacy in prostate cancer except in rare cases with mismatch repair defects.^18–21^ Prostate cancer typically has a myeloid-dominant immune cell infiltrate that largely outnumbers effector lymphocytes and can exert immunosuppression through described mechanisms such as adenosine signaling, IL-23 activation of the androgen receptor, and recruitment of additional suppressive myeloid cells.^22–26^ Regulatory T cells (Tregs) are another immunosuppressive population that can be recruited by and co-occur with suppressive macrophage populations, are associated with worse prognosis, and expand under androgen deprivation.^27–30^ A thorough understanding of tumor-immune system interactions is required to develop better immunotherapy approaches for prostate cancer, as tumor-intrinsic genetic alterations shape the immune microenvironment and drive heterogeneity in immunotherapy response and resistance.^18,31–33^

DDR defects, such as BRCA2 deficiency, cause genomic instability that drives cGAS/STING pathway activation, resulting in an innate immune response and increased interferon (IFN) and cytokine production.^34–38^ By this mechanism, cGAS/STING activation is thought to act as a tumor suppressor, promoting anti-tumor immunity by recruiting and activating immune cells. However, paradoxically, constitutive activation has also been reported to promote tumor progression and immunosuppression, in part through recruitment of immunosuppressive cells and upregulation of checkpoint ligands.^39–42^ Therefore, pathway activation alone does not predict the immune consequences of BRCA2 loss. These immune effects remain poorly characterized in prostate cancer compared to breast or ovarian cancer, where the consequences of BRCA2 loss for tumor development and therapeutic targeting have been more extensively studied.^36,43–46^ The sources of DNA damage and impact of DNA repair defects can vary by tissue type, reflecting differences in metabolism, inflammation, hormone responsiveness, and environmental stressors, necessitating further understanding of tissue-specific biology. How BRCA2 loss shapes the tumor microenvironment (TME) in prostate cancer therefore remains to be defined and is likely distinct from other tumor types. Here, we generated an isogenic, immunocompetent model of *Brca2*-deficient prostate cancer in which the microenvironmental consequences of BRCA2 loss can be dissected and targeted. This identified an interferon-responsive macrophage program associated with enrichment of immunosuppressive CCR8⁺ regulatory T cells, revealing a therapeutically targetable axis.

## Results

### Establishing a syngeneic *Brca2*-deficient mouse model of prostate cancer

To establish a preclinical model of BRCA2-deficient prostate cancer, CRISPR/Cas9 was used to introduce frameshift mutations (*Brca2^fs^*) in the *Brca2* gene in the androgen-dependent MyC-CaP mouse prostate cancer cell line (Fig. 1A).^47^ Isogenic MyC-CaP cell lines with loss of function (LOF) homozygous or compound heterozygous frameshift mutations were obtained and validated by Sanger sequencing and whole exome sequencing (WES) (Extended Data Fig. 1A-B). Frameshift mutations were predicted to generate protein truncation and RNA sequencing (RNA-seq) of *Brca2^fs^* isogenic MyC-CaP clones and RT-qPCR of cell lines and tumors demonstrated a significant reduction in *Brca2* gene expression consistent with nonsense-mediated decay (Fig. 1B-C, Extended Data Fig. 1C).

**Fig. 1:**
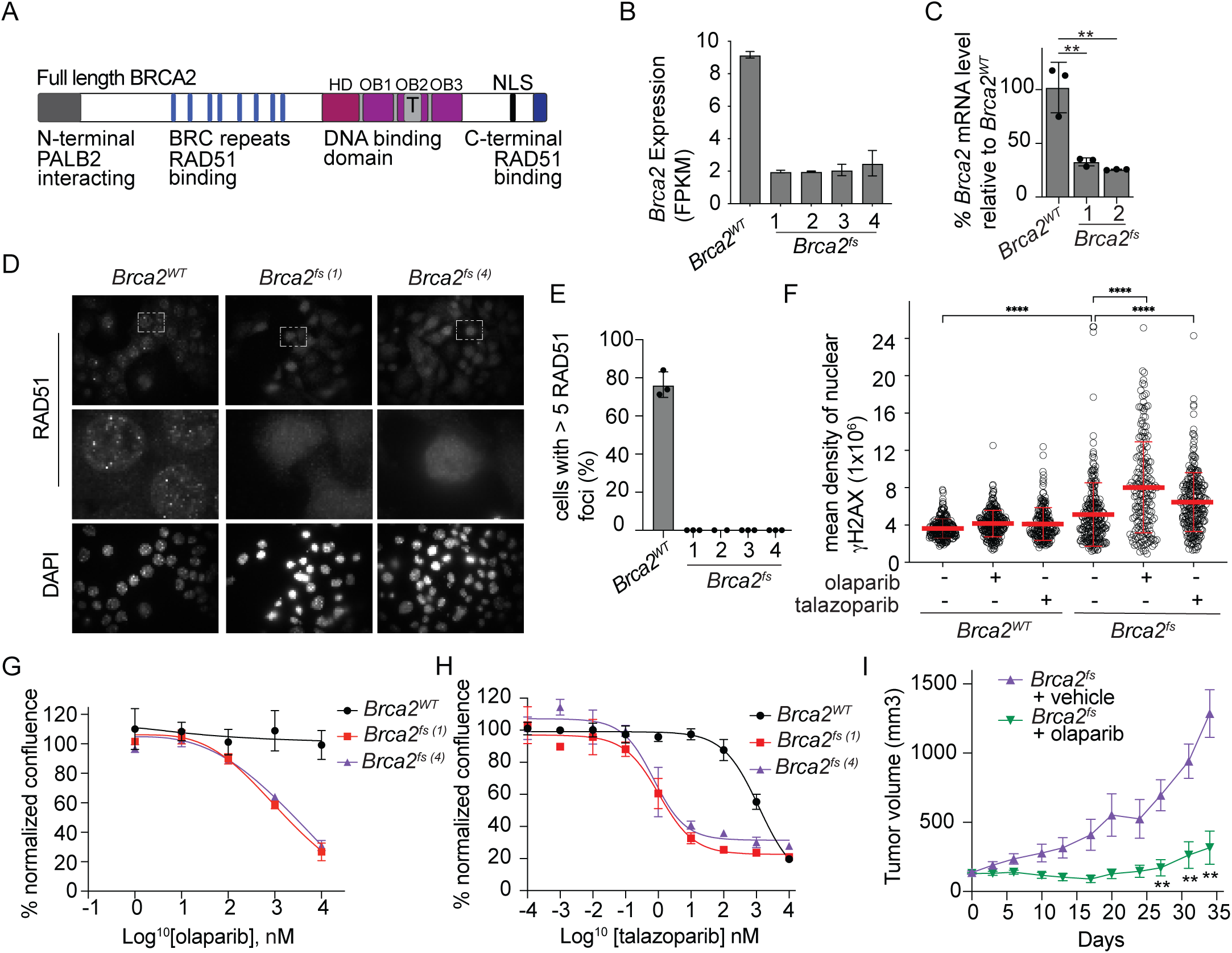
An immunocompetent mouse model of BRCA2-deficient prostate cancer. (A) Schematic of BRCA2 domain structure, with conserved domains with sites of RAD51 interaction indicated. (B) Normalized expression from RNA sequencing (RNA-seq) of *Brca2^fs^* MyC-CaP cell line clones in fragments per kilobase of transcripts per million reads (FPKM). (C) RT-qPCR of *Brca2* expression levels of *Brca2^fs^* MyC-CaP clones relative to isogenic *Brca2^WT^* MyC-CaP cells. (D) Immunofluorescence (IF) imaging of RAD51 foci 24 h after treatment with mitomycin C (MMC). Top row, 63×, and second row enlarged images to demonstrate RAD51 foci in individual cells. (E) Quantification of RAD51 foci 24 h after treatment with MMC in isogenic *Brca2^WT^* MyC-CaP cells and *Brca2^fs^* MyC-CaP clones. (F) Quantification of mean density of nuclear γH2AX by IF in *Brca2^WT^* compared to *Brca2^fs^* that are untreated, after olaparib (2 µM), and after talazoparib (2 nM). (G) Cell confluence analysis by Incucyte after treatment with olaparib comparing *Brca2^WT^* and *Brca2^fs^* MyC-CaP cells. (H) Cell confluence by Incucyte after treatment with talazoparib comparing *Brca2^WT^* and *Brca2^fs^* MyC-CaP cells. (I) FVB mice were injected subcutaneously with *Brca2^fs^* MyC-CaP cells. Mice received either vehicle or olaparib (75 mg/kg, intraperitoneal (IP) daily). Tumor volumes were measured twice weekly and graphed versus day post the start of treatment. Each point shows mean tumor volume and error bars are SEM. n = 13 vehicle and 12 olaparib. Tumor volumes were compared at each timepoint by mixed-effects model (REML) with Bonferroni’s correction for multiple comparisons. Asterisks indicate significance of *Brca2^fs^* olaparib versus Brca*2^fs^* vehicle, otherwise not significant. ** P < 0.01.

To functionally assess *Brca2* LOF, we assessed RAD51 foci levels after DNA damage with mitomycin C (MMC) in several clones (1-4). Consistent with BRCA2 deficiency, *Brca2^fs^* isogenic MyC-CaP clones showed loss of observable RAD51 foci formation (Fig. 1D-E). Immunofluorescence (IF) analysis of the DNA damage marker γH2AX in isogenic cell lines revealed significantly higher levels of γH2AX in *Brca2^fs^* cells compared to *Brca2^WT^* cells which further increased upon treatment with the PARPi olaparib and talazoparib (Fig. 1F). As expected, *Brca2^fs^* MyC-CaP clones showed sensitivity to PARPi compared to isogenic *Brca2^WT^* cells (Fig. 1G-H, Extended Data Fig. 1D-E).^13,48^ *Brca2^fs^* cells were also sensitive to the DNA damaging agents, SN-38 and cisplatin (Extended Data Fig. 1F-G). To test PARPi sensitivity *in vivo*, *Brca2^fs^* cells (clone 1) were implanted subcutaneously into the flanks of immunocompetent FVB/N mice; upon tumors reaching 100-200mm^3^ in size, mice were randomized to daily olaparib (75 mg/kg) or vehicle. Mice treated with olaparib had significantly smaller tumors and longer survival time compared to vehicle-treated animals (Fig. 1I, Extended Data Fig. 1J-K) confirming the expected sensitivity. Together, these data establish an HR-deficient, PARPi-sensitive syngeneic model of *Brca2*-deficient prostate cancer.

### *Brca2* loss promotes increased cGAS-STING pathway activation in prostate cancer cells

Aberrant DNA accumulation in the cell cytoplasm can trigger cGAS/STING pathway activation and promote type I interferon response.^37,38^ BRCA2 deficiency leads to genomic instability and has been linked to cGAS/STING pathway activation in breast and ovarian cancer cells.^44–46^ Consistent with this, *Brca2^fs^* prostate cancer cells showed increased micronuclei formation with cGAS recruitment, by IF, in addition to elevated γH2AX (Fig. 1F, 2A-B). *Brca2^fs^* cells showed increased phosphorylation of core cGAS/STING pathway signaling proteins including IRF3, STING, and TBK1, which was further enhanced by genotoxic treatment with olaparib (Fig. 2C-D). Downstream of cGAS/STING activation, interferon-responsive chemokines *Ccl5* and *Cxcl10* were transcriptionally elevated and further induced by treatment with olaparib in *Brca2^fs^* prostate cancer cells (Fig. 2E-F). Genetic deletion of either *Sting* (*Tmem173)* or *Cgas* eliminated or markedly reduced STING phosphorylation, respectively (Fig. 2G). This suggests that the DNA-damage driven STING activation in *Brca2^fs^* depends on the canonical cGAS/STING pathway.^37,38^ To assess interferon pathway activity in *Brca2^fs^* prostate cancer *in vivo*, bulk RNA-sequencing (RNA-seq) was performed on *Brca2^WT^* and *Brca2^fs^* tumors (n=10, per genotype). Gene set enrichment analysis (GSEA) confirmed a dominant interferon response in *Brca2^fs^* tumors compared to *Brca2^WT^* tumors, with interferon-α and interferon-γ emerging as the top enriched hallmarks (Fig. 2H). Differential gene expression analysis showed a coordinated increase in interferon-stimulated genes (such as *Usp18, Irf7, Irf9, Isg15, Oas* family factors, and others) (Fig. 2I). Together, these data show that *Brca2* deficiency is associated with a robust interferon program *in vivo* and support a cGAS/STING dependent interferon response in *Brca2*-deficient prostate cancer cells.

**Fig. 2:**
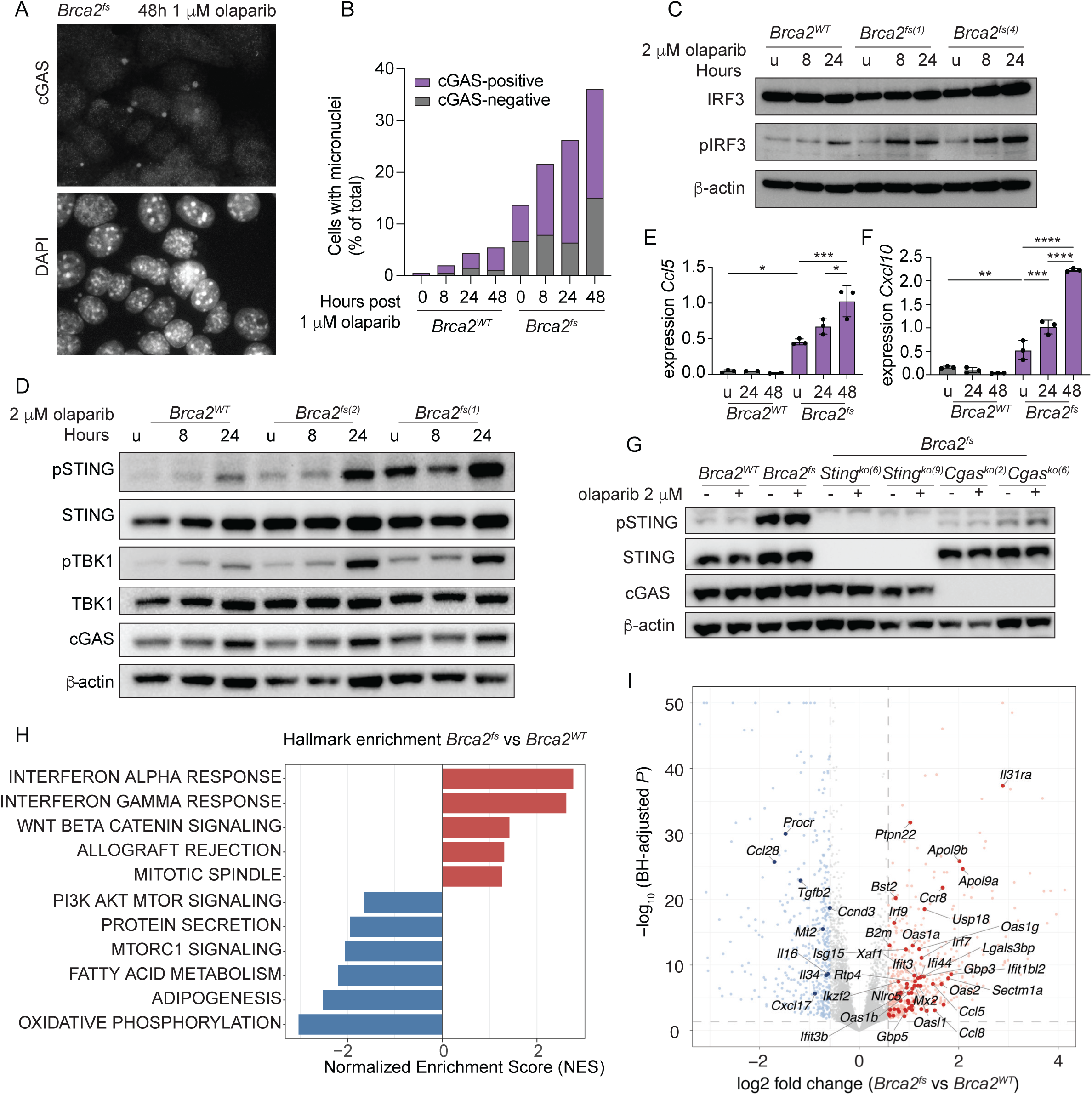
Activation of the innate immune pathway in BRCA2-deficient prostate cancer. (A) IF image representation of cGAS and DAPI costained nuclei in *Brca2^fs^* MyC-CaP cells 48 h post 1 µM olaparib treatment. (B) Quantification of micronuclei formation in *Brca2^WT^* and *Brca2^fs^* cells after 1 µM olaparib treatment as visualized in A. (C) Western blot analysis of IRF3 and phosphorylated IRF3 (pIRF3) of *Brca2^WT^* and *Brca2^fs^* (clones 1, 4) cells untreated (u) or 8 h and 24 h after treatment with 2 µM olaparib. (D) Western blot analysis of STING, phosphorylated STING (pSTING), TBK1, phosphorylated TBK1 (pTBK1), and cGAS of *Brca2^WT^* and *Brca2^fs^* (clones 1, 2) cells untreated (u) or 8 h and 24 h after treatment with 2 µM olaparib. (E-F) RT-qPCR of *Ccl5* and *Cxcl10* mRNA expression levels of *Brca2^WT^* and *Brca2^fs^* MyC-CaP clones either untreated (u) or 24 h and 48 h after 2 µM olaparib treatment. Values are normalized to *Gapdh*. (G) Western blot analysis of STING, pSTING, and cGAS of *Brca2^WT^*, *Brca2^fs^, Brca2^fs^*;*Sting^ko^* and *Brca2^fs^*;*Cgas^ko^* clones untreated (u) and 24 h after treatment with 2 µM olaparib. (H) Hallmark pathway enrichment using fgsea on Wald-statistic ranked-genes from *Brca2^WT^*and *Brca2^fs^* tumors. Bars indicate normalized enrichment score (NES) for the top five up- and six down-regulated pathways in *Brca2^fs^* tumors of Hallmark gene sets reaching fgsea adjusted p < 0.25. Red bars indicate enriched and blue bars indicate depleted pathways in *Brca2^fs^* tumors compared to *Brca2^WT^* tumors. (I) Volcano plot of differentially expressed genes in *Brca2^WT^* and *Brca2^fs^* tumors (DESeq2 Wald test with ashr shrinkage; padj from Benjamini–Hochberg adjustment). Log_2_fold change plotted on the x-axis and −log10(BH-adjusted P) on the y-axis. Significance threshold of adjusted p < 0.05 and |log_2_fold change| >= log2(1.5). Values exceeding the axis limits are plotted at the boundary. Deep red (up) and dark navy (down) points denote significant hallmark genes annotated from MSigDB Hallmark Interferon-α Response, Interferon-γ Response, or Allograft Rejection gene sets. The 25 most significant plus additionally identified mouse-specific ISGs, *Ccr8*, and *Ccl8* are labeled.

### *Brca2* loss is associated with a multicompartment interferon response

We performed single-cell RNA sequencing (scRNA-seq) on six *Brca2^fs^* and *Brca2^WT^* (n=3 per genotype) tumors grown in syngeneic FVB/N mice to define the TME associated with *Brca2* loss and tumor-intrinsic interferon responses. Integrated scRNA-seq analysis resolved 16 transcriptionally distinct cell types spanning tumor, stromal/vascular, and immune compartments (Fig. 3A, Extended Data Fig. 2A). Overall cellular composition was similar across cell types except for a significant increase in cancer-associated fibroblasts (CAF) and T/NK cells in *Brca2^fs^* tumors compared to *Brca2^WT^* (Fig. 3B, Extended Data Fig. 2B). GSEA analysis using the Hallmark gene set collection confirmed the interferon-α response as a top enriched pathway in the tumor cell compartment in *Brca2^fs^* tumor cells, alongside coordinated downregulation of TNFɑ/NF-κΒ and oxidative-phosphorylation pathways. Interferon-α signature scores were correspondingly elevated in *Brca2^fs^*tumor cells (Fig. 3C-D).

**Fig. 3:**
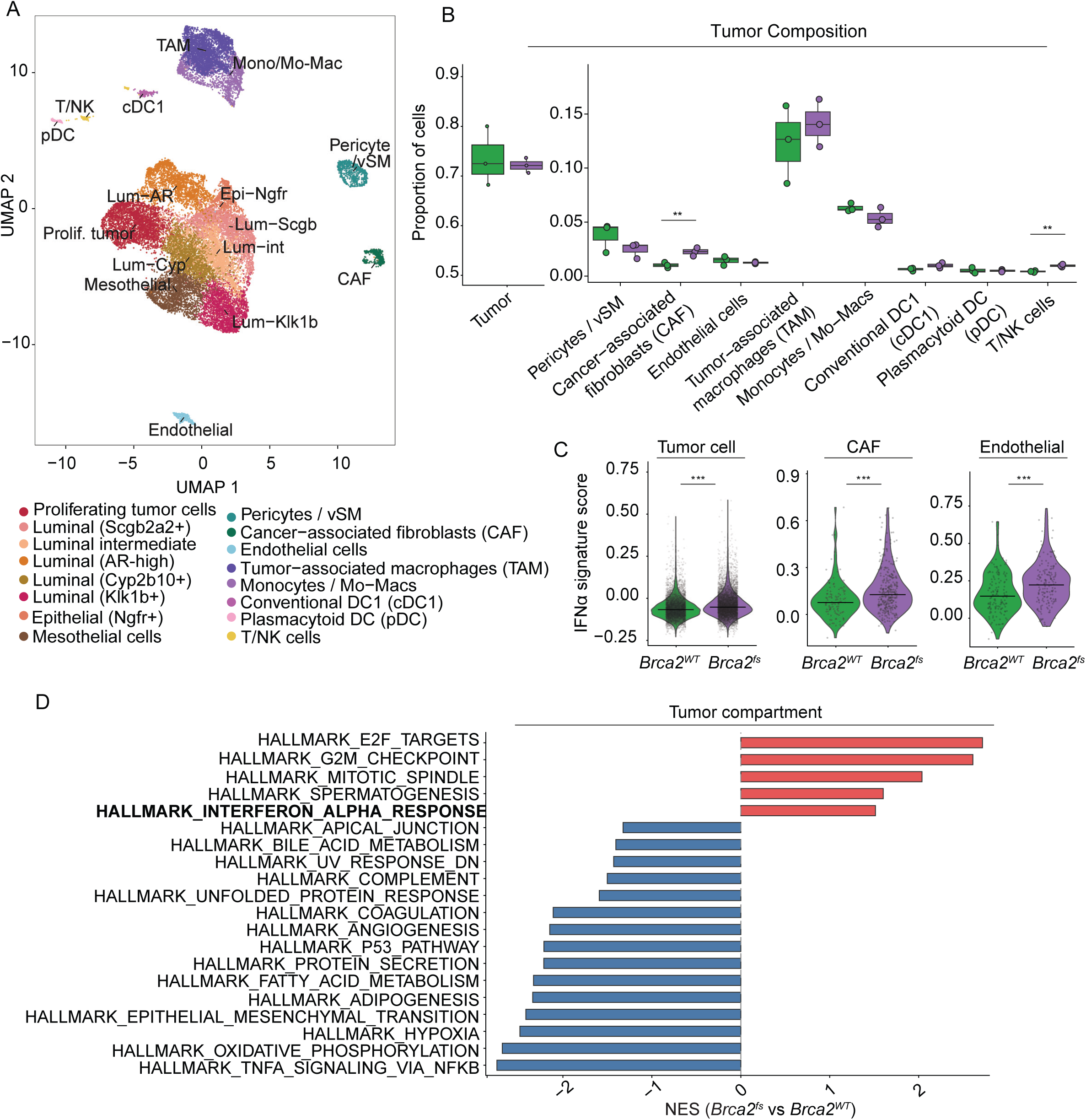
Single-cell RNA sequencing (scRNA-seq) reveals tumor-intrinsic interferon activation. (A) Integrated UMAP of all cells (23,249 cells across the 16 clusters) from *Brca2^WT^* and *Brca2^fs^* tumors (n=3 mice/group). Colored for the 16 annotated cell types detected. (B) Box plot of cell-type composition quantified in *Brca2^WT^* and *Brca2^fs^* tumors as a proportion of all the cells. Tumor/epithelial cell compartment is merged. Points represent individual samples; propeller moderated t-test, ** FDR < 0.01 otherwise nonsignificant. (C) Violin plots demonstrating per-cell distribution of Hallmark IFN-α response signature score (AddModuleScore) of *Brca2^WT^* and *Brca2^fs^* tumors for the tumor cell compartment, cancer-associated fibroblast (CAF), and endothelial cells shown separately. The solid line denotes the median and significance as marked (*** p <0.001). (D) Hallmark gene-set enrichment (GSEA) of the tumor-compartment by pseudobulk transcriptome of *Brca2^fs^ vs Brca2^WT^* tumors. Red bars indicate enriched and blue bars are down-regulated pathways shown by NES (preranked GSEA, padj < 0.05).

Interferon-response signatures were also higher in CAF and endothelial cell compartments of *Brca2^fs^* tumors (Fig. 3C, Extended Data Fig. 2C-E). CAFs are established responders to interferons and interferon-response CAF subsets induced by STING agonism have been described.^49,50^ Expansion of this population in *Brca2^fs^* tumors suggests that tumor-associated interferon may enrich an interferon-responsive fibroblast population. Endothelial cells are similarly responsive, and tumor-derived cGAMP can activate endothelial STING, promoting immune cell infiltration.^51,52^ These findings show that BRCA2-deficient tumors exhibit a multi-compartment interferon response across tumor, fibroblast, and endothelial populations, establishing a microenvironment-wide interferon responsive state.

### An interferon-activated immunosuppressive macrophage compartment in *Brca2*-deficient prostate cancer

After the tumor cell compartment, macrophages constituted the next largest cell population and the largest immune compartment, consistent with the previously described myeloid-predominant tumor immune microenvironment of prostate cancer (Fig. 3B).^31^ To resolve heterogeneity in this compartment, myeloid cells were re-clustered, resolving ten subclusters comprising seven macrophage subtypes, classical monocytes, conventional type-1 dendritic cells (cDC1), and plasmacytoid dendritic cells (pDC) (Fig. 4A, Extended Data Fig. 3A-B). In *Brca2^fs^* tumors, macrophage composition was altered compared to wild-type: resident tumor-associated macrophages (Res-TAM) and perivascular TAM (PV-TAM) were expanded, and lipid-associated macrophages (LAM) and proliferating TAM (Prolif-TAM) were reduced (Fig. 4B).

**Fig. 4:**
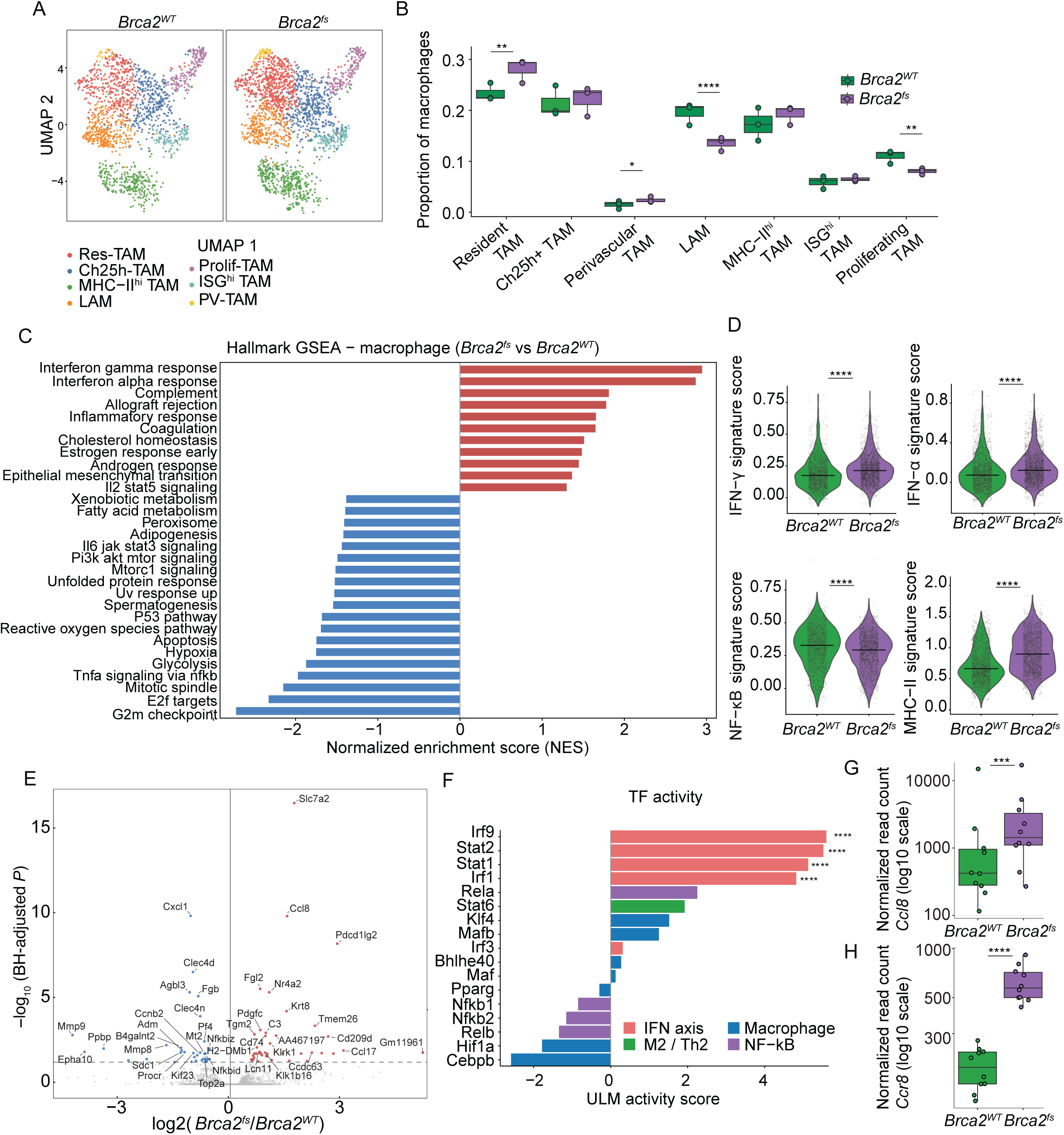
Interferon-associated transcriptional reprogramming in tumor-associated macrophages is accompanied by increased *Ccl8* expression. (A) UMAP of myeloid re-clustering, macrophage clusters only (3,918 cells in seven macrophage clusters, dendritic cells and monocytes excluded), split by *Brca2^WT^* and *Brca2^fs^* tumors. Colored by the 7 macrophage clusters: resident TAM (Res-TAM), Ch25h⁺ TAM (Ch25h-TAM), MHC-II^hi^ TAM, lipid-associated macrophages (LAM), proliferating TAM (Prolif-TAM), ISG^hi^ TAM, and perivascular TAM (PV-TAM). (B) Box plot of macrophage subtype composition in *Brca2^WT^* and *Brca2^fs^* tumors as per-sample proportion of macrophages. Differential abundance by negative-binomial GLM (cell counts ∼ group, log-total offset). (C) Hallmark gene-set enrichment (GSEA) of the macrophage-compartment by pseudobulk transcriptome of *Brca2^fs^ vs Brca2^WT^* tumors. Red bars are significantly up- and blue bars significantly down-regulated pathways shown by NES (preranked GSEA, FDR < 0.05). (D) Violin plots demonstrating per-cell distribution of Hallmark IFN-γ, IFN-α and TNFɑ/NF-κB signatures and an MHC-II signature response score (AddModuleScore) of *Brca2^WT^* and *Brca2^fs^* tumor macrophages. The solid line denotes the median. Significance by two-sided Wilcoxon rank-sum test on per-cell scores, as marked. (E) Volcano plot of differentially expressed genes in *Brca2^fs^ vs Brca2^WT^* macrophage compartment (pseudobulk DESeq2 Wald test). Log2 fold change on the x-axis and -log10(BH-adjusted P) on the y-axis. Red points denote significantly up-regulated genes in *Brca2^fs,^* and blue points genes significantly down-regulated (FDR < 0.05). Labeled points emphasize significant macrophage chemokines, checkpoint-ligand, and antigen-presentation genes. (F) Transcription-factor activity inference in the macrophage compartment in *Brca2^fs^ vs Brca2^WT^* tumors (decoupleR, CollecTRI regulons; univariate linear model (ULM)) for candidate transcription factors. Activity was scored by ULM on the macrophage-compartment pseudobulk DESeq2 Wald statistic, and bars are colored by the functional family. Significance of the ULM activity score is marked, with the Benjamini–Hochberg adjustment taken across all tested regulons. (G-H) Box plot of DESeq2-normalized bulk RNA-seq read counts (log10 scale) for *Ccl8* and *Ccr8* from *Brca2^WT^* and *Brca2^fs^* tumors. Each point represents a tumor; differential expression by DESeq2 Wald test with ashr shrinkage. Multiple comparisons were corrected by the Benjamini–Hochberg procedure throughout; *FDR < 0.05, **FDR < 0.01, ***FDR < 0.001, ****FDR < 0.0001.

Hallmark GSEA of the macrophage compartment identified interferon-α and interferon-γ as top enriched pathways in *Brca2^fs^* tumors, alongside enrichment of complement, allograft rejection, inflammatory response, and IL2/STAT5 signaling, with concurrent downregulation of TNFɑ/NF-κΒ signaling (Fig. 4C). Signature scoring reproduced this pattern and additionally showed elevated MHC II signature (Fig. 4D). The interferon-activated macrophage phenotype occurred without a classical M1/M2 polarization (Extended Data Fig. 3F). In *Brca2^fs^* tumors, differential gene expression analysis confirmed gene level upregulation of MHC-II antigen-presentation genes (*Cd74, H2-DMb1*), and induction of immunosuppressive mediators: Treg recruiting chemokines *Ccl8* (CCR8 ligand) and *Ccl17* (CCR4 ligand), inhibitory checkpoint ligand *Pdcd1lg2* (PD-L2), *Fgl2,* arginine transporter *Slc7a2,* and NF-κB repressor *Nr4a2/3* (Fig. 4E). Inferred transcription-factor activity analysis (decoupleR, CollecTRI mouse regulons) identified the interferon-stimulated-gene-factor-3 (ISGF3) components *Irf9*, *Stat2*, and *Stat1*, together with Irf1, as the most strongly activated transcription factors of candidate transcription factors examined in *Brca2^fs^* tumors relative to *Brca2^WT^* (Fig. 4F). In contrast, NF-κB-family factors did not have significantly increased activation, supporting predominantly canonical interferon (ISGF3/IRF1) as the pathway most strongly associated with the macrophage program, rather than NF-κB signaling. *Ccl8*, which encodes a Treg-recruiting chemokine and CCR8 ligand,^53–55^ was among the most significantly upregulated genes in the macrophage compartment of *Brca2^fs^* tumors (Fig. 4E). *Ccl8* expression was induced across macrophage clusters (Extended Data Fig. 3C-E) and correlated positively with the interferon-stimulated-gene signature and interferon transcription factors (*Stat1, Stat2, Irf7*), and inversely with NF-κB factors (*Rela, Relb, Nfkb1/2*) (Extended Data Fig. 3G).

Given the significant induction of *Ccl8* and other Treg-recruiting chemokines in *Brca2^fs^* tumors, we focused on the CCL8-CCR8 axis, a known chemokine-receptor pathway capable of recruiting immunosuppressive CCR8⁺ T cells.^53–55^ Low T cell numbers detected by scRNA-seq precluded statistical analysis of CCR8⁺ Tregs; however, bulk RNA-seq of *Brca2^fs^* and *Brca2^WT^* tumors further confirmed significantly increased expression of both *Ccl8* and *Ccr8* (Fig. 4G-H). These data support the presence of an interferon-associated, immunosuppressive CCL8-CCR8 axis in *Brca2*-deficient tumors.

### CCR8⁺ regulatory T cells are enriched in *Brca2*-deficient tumors and are therapeutically targetable

To evaluate whether *Ccl8* upregulation in *Brca2*-deficient tumors is associated with increased tumor-infiltrating Tregs, T cell populations were quantified by spectral flow cytometry of *Brca2^WT^* and *Brca2^fs^* tumors (Extended Data Fig. 4A). CD8⁺ T cell abundance did not differ significantly; however, *Brca2^fs^* tumors had significantly increased proportions of CD4⁺ T cells and CD4⁺CD25⁺FOXP3⁺ Tregs (Fig. 5A-C). In line with transcriptional upregulation of the *Ccl8*–*Ccr8* axis (Fig. 4G-H), CCR8⁺ Tregs (CD4⁺CD25⁺FOXP3⁺CCR8⁺) were significantly elevated in *Brca2^fs^* tumors compared to *Brca2^WT^* tumors (Fig. 5D-E). These data establish enrichment of CCR8⁺ Tregs in *Brca2*-deficient tumors, consistent with a more immunosuppressive TME.

**Fig. 5:**
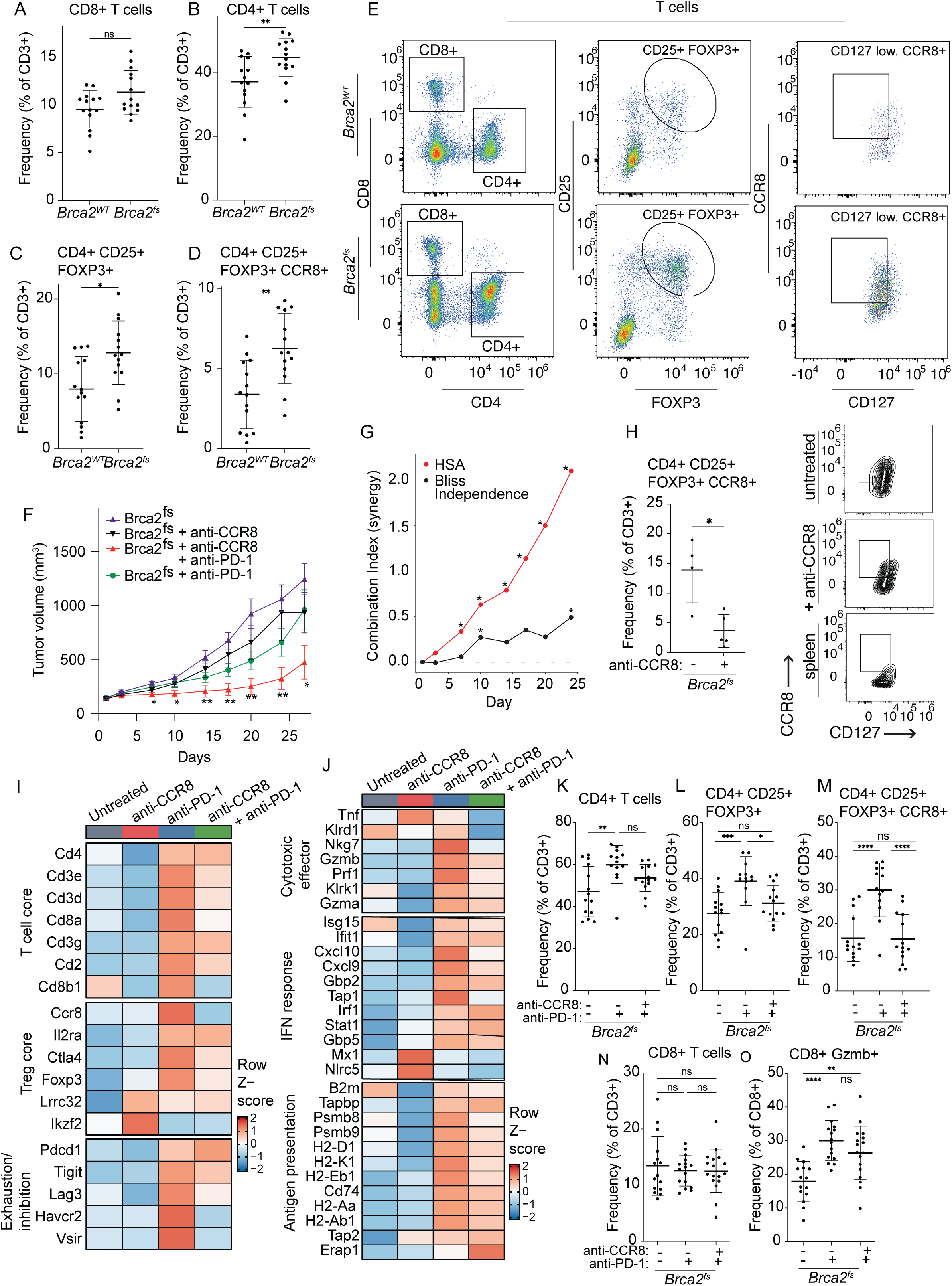
Combination of anti-PD-1 and anti-CCR8 retains effector recruitment while eliminating PD-1-induced CCR8⁺ Treg expansion. (A-D) Spectral flow cytometry quantification of T-cell subsets in *Brca2^WT^* and *Brca2^fs^* tumors. CD8⁺ T cells, CD4⁺ T cells, regulatory T-cell subset (CD4⁺ CD25⁺ FOXP3⁺), and CCR8⁺ regulatory T-cell subset (CD4⁺ CD25⁺ FOXP3⁺ CCR8⁺) as a percent of the CD3⁺ population. Each point represents one tumor, and lines indicate mean ± SD; n = 14 per genotype, pooled from two independent experiments stained and acquired with the same panel. Two-tailed Mann-Whitney test. *P < 0.05, **P < 0.01; ns, not significant. (E) Representative spectral flow cytometry plots from *Brca2^WT^* and *Brca2^fs^* tumors used for quantification in A-D. (F) Male FVB mice were injected subcutaneously with *Brca2^fs^* MyC-CaP cells. Mice received either no treatment (n=8), anti-CCR8 monoclonal antibody (mAb) alone (10 mg/kg, i.p. three days a week for 8 doses, n=9), anti-PD-1 mAb alone (10 mg/kg, i.p. three days a week for 8 doses, n=10), or combination anti-CCR8 and anti-PD-1 mAb (same dose and schedule, n=9). Each point shows mean tumor volume and error bars are SEM. Tumor volumes were compared at each timepoint by mixed-effects model (REML) with Tukey’s correction for multiple comparisons. Asterisks indicate significance versus the untreated *Brca2^fs^*, otherwise not significant. *P < 0.05, **P < 0.01. (G) Combination index calculated for anti-CCR8 and anti-PD-1 in F. Calculated by Highest Single Agent (HSA, in red) and Bliss Independence (black) models from tumor growth through day 24. Values > 0 indicate synergy and asterisks indicate significant synergy by one-sided t-test on per mouse relative tumor volume. *P < 0.05, **P < 0.01, ***P < 0.001. (H) Left: spectral flow cytometry quantification of CCR8⁺ regulatory T-cell subset (CD4⁺ CD25⁺ FOXP3⁺ CCR8⁺) as a percent of the CD3⁺ population in *Brca2^fs^* tumors from mice either untreated or treated with anti-CCR8 mAb. Each point represents one tumor, and lines indicate mean ± SD; n = 4 untreated and 5 anti-CCR8. Two-tailed unpaired t test with Welch’s correction. *P < 0.05. Right: representative spectral flow cytometry plots of CCR8⁺ regulatory T-cell subset (CD4⁺ CD25⁺ FOXP3⁺ CCR8⁺) from untreated *Brca2^fs^* tumor, anti-CCR8 mAb treated *Brca2^fs^* tumor, and untreated spleen. (I-J) Group mean heatmap of (I) T cell and regulatory T cell related genes (T cell core, Treg core, Exhaustion/inhibition) and (J) cytotoxic effector, IFN response, and antigen-presentation signature genes across treatment arms (untreated, anti-CCR8 mAb, anti-PD-1 mAb, and anti-CCR8 + anti-PD-1 mAb). Bulk-tumor RNA-seq counts were variance-stabilized (DESeq2 VST), averaged per arm, and row-normalized to Z-scores; red indicates higher expression and blue indicates lower expression relative to the crossarm mean. n = 7 untreated, 7 anti-CCR8, 8 anti-PD-1 and 8 anti-CCR8 + anti-PD-1 tumors, from a separate cohort treated as in F but for 6 doses. (K-O) Spectral flow cytometry quantification of T-cell subsets: CD4⁺ T cells, regulatory T-cell subset (CD4⁺ CD25⁺ FOXP3⁺), CCR8⁺ regulatory T-cell subset (CD4⁺ CD25⁺ FOXP3⁺ CCR8⁺), CD8⁺ T cells as a percent of the CD3⁺ population, and activated CD8⁺ T cells (granzyme B (Gzmb⁺)) as a percent of CD8⁺ cells in *Brca2^fs^* tumors: untreated, after anti-PD-1 mAb, or anti-CCR8 + anti-PD-1 mAb. Each point represents one tumor, and lines indicate mean ± SD. K-M: n = 14, 13, 14; N-O: n = 15, 16, 17; each pooled from two independent experiments. Ordinary one-way ANOVA with Tukey’s correction for multiple comparisons. *P < 0.05, **P < 0.01, ***P < 0.001, ****P < 0.0001; ns, not significant.

Given this enrichment of CCR8⁺ Tregs, together with induction of PD-1 ligands (Extended Data Fig. 3E), we tested whether targeting CCR8⁺ Tregs for depletion in *Brca2^fs^* prostate tumors, alone or combined with anti-PD-1 monoclonal antibody (mAb) checkpoint blockade, could enhance anti-tumor immunity. *Brca2^fs^* tumors were established in FVB/N mice, which were randomized to receive anti-CCR8, anti-PD-1, or the combination of the two mAbs (Fig. 5F). Combined anti-CCR8 and anti-PD-1 treatment strongly suppressed tumor growth, while the individual mAbs did not have a significant effect. The combined effect exceeded both the better single agent and predicted additive activity, indicating synergy by HSA (Highest Single Agent) and Bliss independence analysis, respectively (Fig. 5G).^56^ Anti-CCR8 mAb effectively depleted CCR8⁺ Tregs (Fig. 5H).

To understand the transcriptional response to each therapy, bulk RNA-seq was performed on *Brca2^fs^* tumors from mice receiving anti-CCR8, anti-PD-1, combined anti-CCR8 and anti-PD-1, or no treatment. Anti-CCR8 monotherapy produced no coordinated immune program changes (Fig. 5I–J, Extended Data Fig. 4B-C). Anti-PD-1 mAb therapy induced an inflammatory response (Extended Data Fig. 4B-C), with coordinated increases in T cell infiltration (*Cd3d/e/g, Cd4, Cd8a*), cytotoxic-effectors (*Gzmb, Prf1, Nkg7*), interferon effectors (*Cxcl9, Cxcl10, Stat1, Irf1*), and antigen-presentation machinery (*B2m, Tap2, Cd74, H2* genes) (Fig. 5I-J). Surprisingly, anti-PD-1 mAb therapy also induced a counter-regulatory immunosuppressive program, further upregulating Treg core genes (*Il2ra, Ctla4, Foxp3*), *Ccr8,* and checkpoint/exhaustion markers (*Pdcd1, Tigit, Lag3, Havcr2, Vsir*) (Fig. 5I-J, Extended Data Fig. 4B-C). Spectral flow cytometry confirmed significant CD4⁺ T cell expansion, with a striking increase in CCR8⁺ Tregs in *Brca2^fs^* tumors treated with anti-PD-1 monotherapy (Fig. 5K-M). Anti-PD-1 therapy elicits a significant inflammatory response in *Brca2^fs^* tumors, but the concurrent expansion of immunosuppressive CCR8⁺ Tregs provides a potential mechanism of treatment failure to checkpoint blockade monotherapy. Combination anti-CCR8 and anti-PD-1 treatment retained elevated interferon response and antigen-presentation programs while depleting CCR8⁺ Tregs (Fig. 5I-J, 5M, Extended Data Fig. 4B-C). Although CD8+ T cell numbers remained unchanged, granzyme B+ (Gzmb+) cytotoxic CD8+ T cells were significantly increased over untreated tumors by both anti-PD-1 monotherapy and the combination (Fig. 5N-O), showing that the effector function induced by checkpoint blockade is retained when CCR8⁺ Tregs are depleted. These data identify CCR8⁺ Tregs as an immunosuppressive barrier in *Brca2*-deficient tumors that is paradoxically expanded by PD-1 blockade and show that their depletion can overcome resistance to PD-1 blockade in this model.

### CCR8⁺ Treg depletion synergizes with PARP inhibition through a distinct mechanism

As PARP inhibition induces both DNA damage and cGAS/STING-dependent immune activation (Fig. 1F, 2),^57,58^ we assessed whether combining PARPi talazoparib with CCR8+ Treg depletion could enhance anti-tumor activity. *Brca2^fs^* tumors were established in FVB/N mice, which were randomized to receive no treatment, low dose talazoparib (0.11 mg/kg, Extended Data Fig. 4D), anti-CCR8 mAb, or the combination of talazoparib and anti-CCR8 mAb. Combined therapy with anti-CCR8 mAb and talazoparib synergistically suppressed tumor growth beyond either monotherapy (Fig. 6A-B). However, spectral flow analysis of treated *Brca2^fs^* tumors suggested that the underlying mechanism differs from that of the anti-PD-1 combination as combination talazoparib with anti-CCR8 mAb significantly increased tumor-infiltrating CD8⁺ T cells (Fig. 6C). Talazoparib monotherapy did not lead to enrichment of CD4⁺ subsets including CCR8⁺ Tregs, in contrast to anti-PD-1 treatment (Fig. 6D). The anti-CCR8-containing combination significantly depleted CCR8⁺ Tregs (Fig. 6E). These data demonstrate that CCR8⁺ Treg depletion synergizes with both immune-checkpoint blockade and PARP inhibition in *Brca2*-deficient prostate cancer, enhancing the immune response through distinct but complementary mechanisms.

**Fig. 6:**
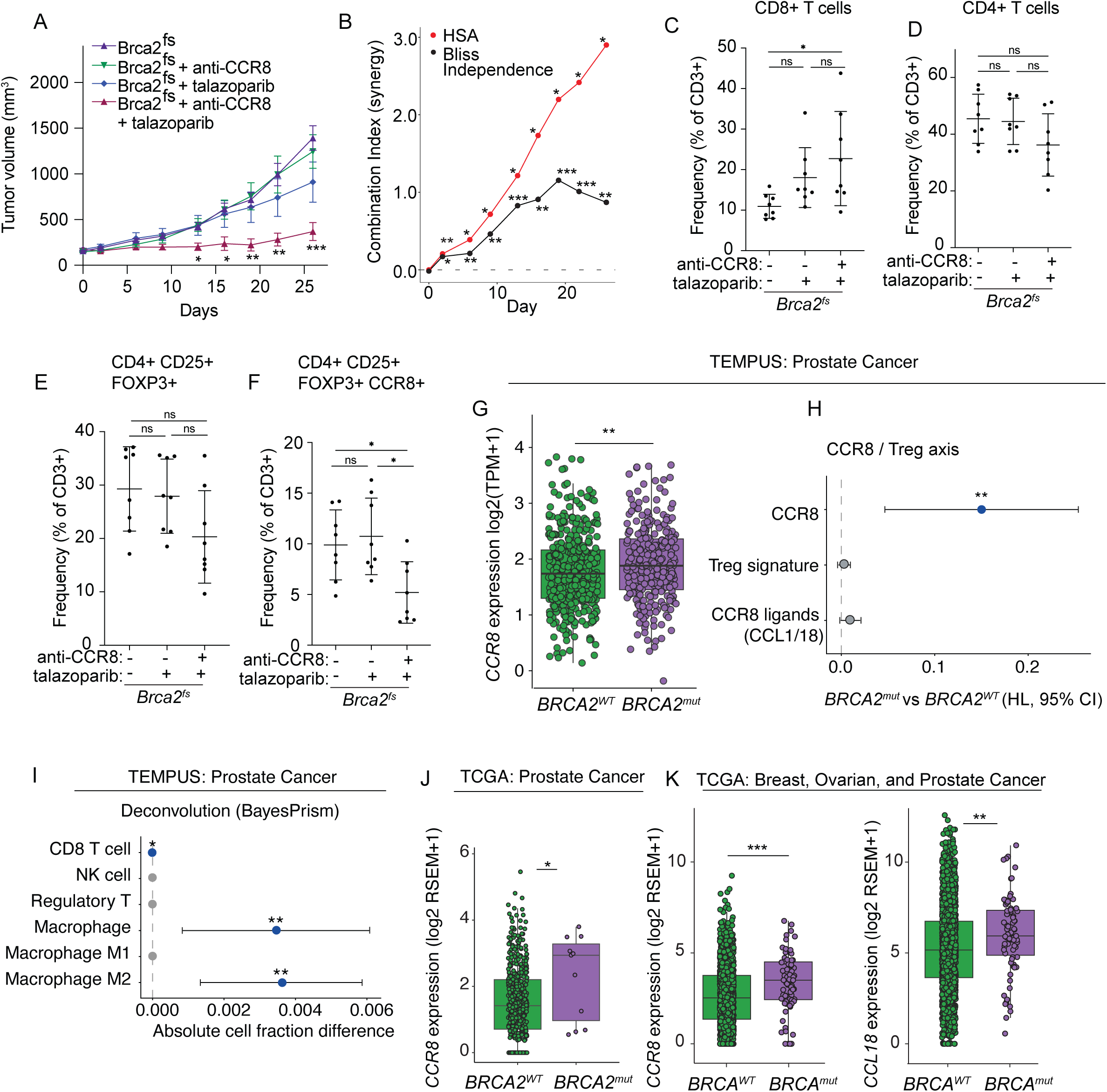
Synergistic activity of CCR8⁺ regulatory-T-cell depletion and PARP inhibition. (A) Male FVB mice were injected subcutaneously with *Brca2^fs^* MyC-CaP cells. Mice received either no treatment (n=10), anti-CCR8 monoclonal antibody (mAb) alone (10 mg/kg, i.p. three days a week for 8 doses, n=10), talazoparib alone (0.11 mg/kg, p.o. five days a week, n=5), or combination anti-CCR8 and talazoparib (same dose and schedule, n=5). Each point shows mean tumor volume and error bars are SEM. Tumor volumes were compared at each timepoint by mixed-effects model (REML) with Tukey’s correction for multiple comparisons. Asterisks indicate significance versus the untreated *Brca2^fs^*, otherwise not significant. *P < 0.05, **P < 0.01, ***P < 0.001. (B) Combination index calculated for anti-CCR8 and talazoparib in A. Calculated by Highest Single Agent (HSA, in red) and Bliss Independence (black) models from tumor growth through day 26. Values > 0 indicate synergy and asterisks indicate significant synergy by one-sided t-test on per mouse relative tumor volume. *P < 0.05, **P < 0.01, ***P < 0.001. (C-F) Spectral flow cytometry quantification of CD8⁺ T cells, CD4⁺ T cells, regulatory T cells (CD4⁺ CD25⁺ FOXP3⁺), and CCR8⁺ regulatory T cells (CD4⁺ CD25⁺ FOXP3⁺ CCR8⁺) as a percent of the CD3⁺ population, from *Brca2^fs^* tumors: untreated, after talazoparib, or anti-CCR8 + talazoparib. Each point represents one tumor, and lines indicate mean ± SD. n = 8 tumors per arm. Ordinary one-way ANOVA with Tukey’s correction for multiple comparisons. *P < 0.05; ns, not significant. (G) CCR8 expression (batch-corrected log2(TPM+1)) in *BRCA2*-mutated (n = 271) versus *BRCA2* wild-type (n = 417) patient prostate cancer biopsy from Tempus AI data set. *BRCA2*-mutated = pathogenic or likely pathogenic short variant or homozygous deletion; *BRCA2* wild-type = no pathogenic alteration in any defined homologous-recombination-repair genes. Wilcoxon rank-sum test (p = 0.005). (H) CCR8-axis modules in *BRCA2*-mutated versus *BRCA2* wild-type tumors (Tempus AI) as Hodges–Lehmann shifts (*BRCA2*-mutated - *BRCA2* wild-type) with 95%CI: CCR8 (the receptor gene), the Treg-context signature, and CCR8 ligands (CCL1, CCL18). Nominal P: *P < 0.05, **P < 0.01, ***P < 0.001. (I) Immune-cell composition by BayesPrism deconvolution, *BRCA2*-mutated vs *BRCA2* wild-type (Tempus AI), as the absolute cell-fraction difference with 95% CI for CD8+ T cells, NK cells, regulatory T cells, total macrophages, and M1 and M2 macrophages; nominal P as in (H). (J) *CCR8* expression (log2 RSEM+1) by *BRCA2* mutation status in TCGA prostate adenocarcinoma (cBioPortal; *BRCA2*-mutated vs *BRCA2* wild-type); two-sided Wilcoxon rank-sum, p = 0.047. (K) *CCR8* (left) and the Treg-recruiting chemokine *CCL18* (right) expression (log2 RSEM+1) by *BRCA1/2* mutation status across combined TCGA breast, ovarian and prostate cohorts (*BRCA*1- or *BRCA2*-mutated vs wild-type); two-sided Wilcoxon rank-sum.

### CCR8⁺ Treg enrichment is conserved in human BRCA2-deficient cancers

We analyzed *BRCA2*-mutated human tumors in the Tempus AI database to assess *CCR8*-axis enrichment in patient specimens. We identified *BRCA2*-mutated (n = 271), and *BRCA2* wild-type (n = 417) primary prostate cancers for analysis. Wild-type was defined as free of pathogenic or likely pathogenic alterations in any homologous recombination repair (HRR genes) and mutant was defined as having pathogenic or likely pathogenic alterations in *BRCA2*. *CCR8* expression was significantly higher in *BRCA2*-mutated tumors compared to wild-type tumors (Fig. 6G) and we saw the same effect on whole transcriptome module scoring, where the Treg signature and CCR8 ligands (*CCL1/CCL18*) shifted in the same direction without reaching significance (Fig. 6H). Consistent with the presence of immunosuppressive myeloid programs, transcriptional immune deconvolution analysis estimated higher total and M2 (immunosuppressive) macrophage fractions in *BRCA2*-mutated prostate cancers, with a small increase in CD8+ T cells, while other immune cell populations were similar to wild-type tumors (Fig. 6I). The *CCR8* association was also observed in *BRCA2*-mutated tumors from the TCGA prostate adenocarcinoma dataset (Fig. 6J) and extended across *BRCA1/2*-mutated breast, ovarian, and prostate cancers, where both *CCR8* and *CCL18* were elevated (Fig. 6K). Together, these data provide population-level, cross-cohort support for the enrichment of the *CCR8*-associated immune program identified in our *Brca2*-deficient mouse model.

## Discussion

Here, we generated an immunocompetent *Brca2*-deficient prostate cancer model and established that loss of BRCA2 function produces a distinct tumor immune microenvironment. Gene expression analysis of *Brca2*-deficient tumors demonstrated an interferon response across cancer cell, fibroblast, and endothelial compartments, an interferon-activated TAM program with upregulation of the CCR8 ligand and Treg chemokine CCL8, and enrichment of highly immunosuppressive CCR8⁺ Tregs. These observations suggested that CCR8⁺ Tregs constrain anti-tumor immunity in *Brca2*-deficient prostate cancer. Accordingly, depletion of CCR8⁺ Tregs with an anti-CCR8 mAb synergized with anti-PD-1 therapy, whereas anti-PD-1 alone paradoxically expanded CCR8⁺ Tregs and induced markers of T cell exhaustion. Our data suggest that this is a targetable mechanism of immune evasion in *BRCA2*-mutated prostate cancer.

A key question in *BRCA2*-mutated prostate cancer is how defective DNA repair alters the tumor immune microenvironment. In *Brca2*-deficient prostate cancer, interferon signaling was evident even in the absence of exogenous genotoxic stress, suggesting that intrinsic DNA damage associated with BRCA2 loss sustains interferon signaling and may shape the surrounding microenvironment during tumorigenesis. The tumor suppressive role of the cGAS/STING pathway underlies efforts to trigger high-intensity STING signaling with radiation, chemotherapy, STING agonists, or PARPi/immunotherapy combinations.^40^ Paradoxically, however, some studies suggest that sustained cGAS/STING activation may promote tumor progression, treatment resistance, and an immunosuppressive TME, highlighting the importance of biological context and the intensity and duration of signaling.^39,40,59^ In *Brca2*-deficient prostate tumors, we found that a constitutive interferon response across the TME was associated with immunosuppression rather than effective anti-tumor immunity.

Our data support a model in which BRCA2 loss generates nucleic acid intermediates that activate cGAS/STING signaling and elicit a tumor-intrinsic interferon response. This mechanism has been described in breast and ovarian models;^44,60^ the present findings extend it to prostate cancer. The innate immune response is further increased by treatment with PARPi, reflecting greater DNA damage and cGAS/STING pathway engagement, consistent with evidence that PARP inhibitors potentiate immune responses in HR-deficient cancers.^57,58^

Interferon signaling was detected not only in tumor cells, but across stromal and immune cell compartments. Consistent with the myeloid predominance in prostate cancer,^22^ macrophages were the largest immune cell population detected, and those in *Brca2*-deficient tumors adopted an interferon-responsive state characterized by increased ISGF3 (*Stat1*, *Stat2*, *Irf9*) and *Irf1* activity without coordinated *NF-κB* pathway activation. This interferon-responsive macrophage state exhibited a significant increase in MHC-II antigen presentation machinery (*Cd74*, *H2-DMb1*) with simultaneous induction of immunosuppressive mediators, including those of immune checkpoints (*Pdcd1lg2*), Treg recruiting chemokines (*Ccl8*, *Ccl17*), *Fgl2,* anti-inflammatory transcription factors *Nr4a2/3,* and the arginine transporter *Slc7a2*.

Of these TAM-derived mediators, *Ccl8* was one of the most significantly upregulated in *Brca2*-deficient tumors and was of particular interest as the ligand for CCR8, a receptor that is selectively expressed on tumor-infiltrating Tregs.^54,55^ In humans, the CCR8 ligands include CCL1 and CCL18,^61^ while in mice CCL8 is the ligand for the CCR8 receptor.^53,62^ In human Tregs, CCL1 induces CCR8-dependent migration that is abolished by a receptor antagonist.^63^ Additionally, murine studies have demonstrated that CCL8 is a potent agonist of CCR8 and promotes T cell recruitment.^53^ CCR8⁺ Tregs are highly immunosuppressive and express multiple immune checkpoints;^54,55^ they have been reported to suppress T cell cytotoxicity, and their density is inversely correlated with granzyme B positivity of CD8+ T cells.^64–66^ Furthermore, high intratumoral *CCR8* expression correlates with poor prognosis in breast, colon, and lung cancers.^54,55,64–66^ *Brca2*-deficient tumors demonstrated a significant increase in *Ccr8* expression and enrichment of CCR8⁺ Tregs, consistent with macrophage *Ccl8* upregulation. These data suggest that in the setting of BRCA2 loss, a sustained interferon response may establish an immunosuppressive circuit in which activated macrophages promote the recruitment of CCR8⁺ Tregs. In a large-scale paired genomic and transcriptomic analysis of human *BRCA2*-mutated prostate cancer from Tempus AI, *CCR8* expression was increased, and deconvolution analysis suggested a rise in total and M2 macrophages. The CCR8 enrichment extended to TCGA *BRCA*1/2-mutated prostate, breast, and ovarian cancers suggesting that CCR8⁺ Tregs enrichment may extend to other HR deficient tumors. These results provide a rationale for targeting the CCR8 axis to overcome this immunosuppressive mechanism and evaluating CCR8-directed therapy in *BRCA2*-mutated prostate cancer.

We targeted CCR8⁺ Tregs *in vivo*, alone or in combination with PD-1 blockade. Neither CCR8⁺ Treg depletion nor PD-1 blockade alone significantly controlled tumor growth. Interestingly, while CCR8 depletion did not promote significant transcriptional changes in *Brca2*-deficient tumors, PD-1 blockade induced a broad inflammatory response including upregulation of inhibitory checkpoints and a compensatory increase in tumor-infiltrating Tregs, most strikingly CCR8⁺ Tregs. Enrichment of CCR8⁺ Tregs in *Brca2*-deficient tumors and their expansion upon anti-PD-1 therapy suggest activity through the CCR8 axis is a primary resistance mechanism that suppresses an effective anti-tumor response to checkpoint blockade. In support of this, the addition of a CCR8 depleting mAb to counter this CCR8⁺ Treg expansion acted synergistically with anti-PD-1 therapy to significantly suppress tumor growth. This is consistent with reported general expansion of Tregs following checkpoint blockade as a mechanism limiting therapeutic efficacy,^67–69^ although CCR8⁺ Treg expansion specifically has not been previously described.

Enrichment of *CCR8* in human *BRCA2*-altered prostate cancer supports the clinical relevance of this axis. Early-phase anti-CCR8 antibody trials are underway in solid tumors, but none include prostate cancer. CCR8 is an attractive therapeutic target because its expression is largely restricted to tumor-resident Tregs across human solid tumors.^54,55^ This selectivity may allow depleting antibodies to spare peripheral Tregs and effector T cells,^70^ potentially uncoupling tumor activity from autoimmune toxicity.^71^ CCR8⁺ Treg depletion could be partnered with either checkpoint inhibition or PARP inhibition, which is already approved in *BRCA2*-mutated prostate cancer. Anti-CCR8 combined with talazoparib showed synergy in controlling tumor growth; however, unlike anti-PD-1, talazoparib did not expand CCR8⁺ Tregs, suggesting a complementary rather than compensatory mechanism.

Tumor genetics drive differences in the TME within prostate cancer, and immunotherapy strategies need to be tailored accordingly.^18,31–33^ Mismatch repair deficiency (MMRd) confers sensitivity to immunotherapy through high tumor mutational burden, and other DDR alterations were initially expected to behave similarly. However, non-MMRd DDR-deficient tumors have not shown comparable immunotherapy response, and their immune consequences are not uniformly activating.^19^ We identify CCR8⁺ Tregs as a barrier to checkpoint blockade efficacy in *Brca2*-deficient prostate cancer and show that their depletion overcomes resistance to anti-PD-1 therapy (Fig. 7). Chronic interferon signaling may promote the immunosuppressive, *Ccl8*-expressing TAM populations and CCR8⁺ Treg recruitment, suggesting that targeting myeloid cells could provide a second point of intervention to disrupt this axis. Therapeutic development may therefore require both relieving immunosuppression and enhancing immune activation. The immune consequences of DDR deficiency likely depend on the specific repair pathway lost, and the broader applicability of this approach will depend on the extent to which these immunosuppressive barriers are shared across DDR-deficient prostate cancer. Defining the immune landscape of prostate cancer across distinct genomic subtypes will be essential to guide rational immunotherapy development.

**Fig. 7:**
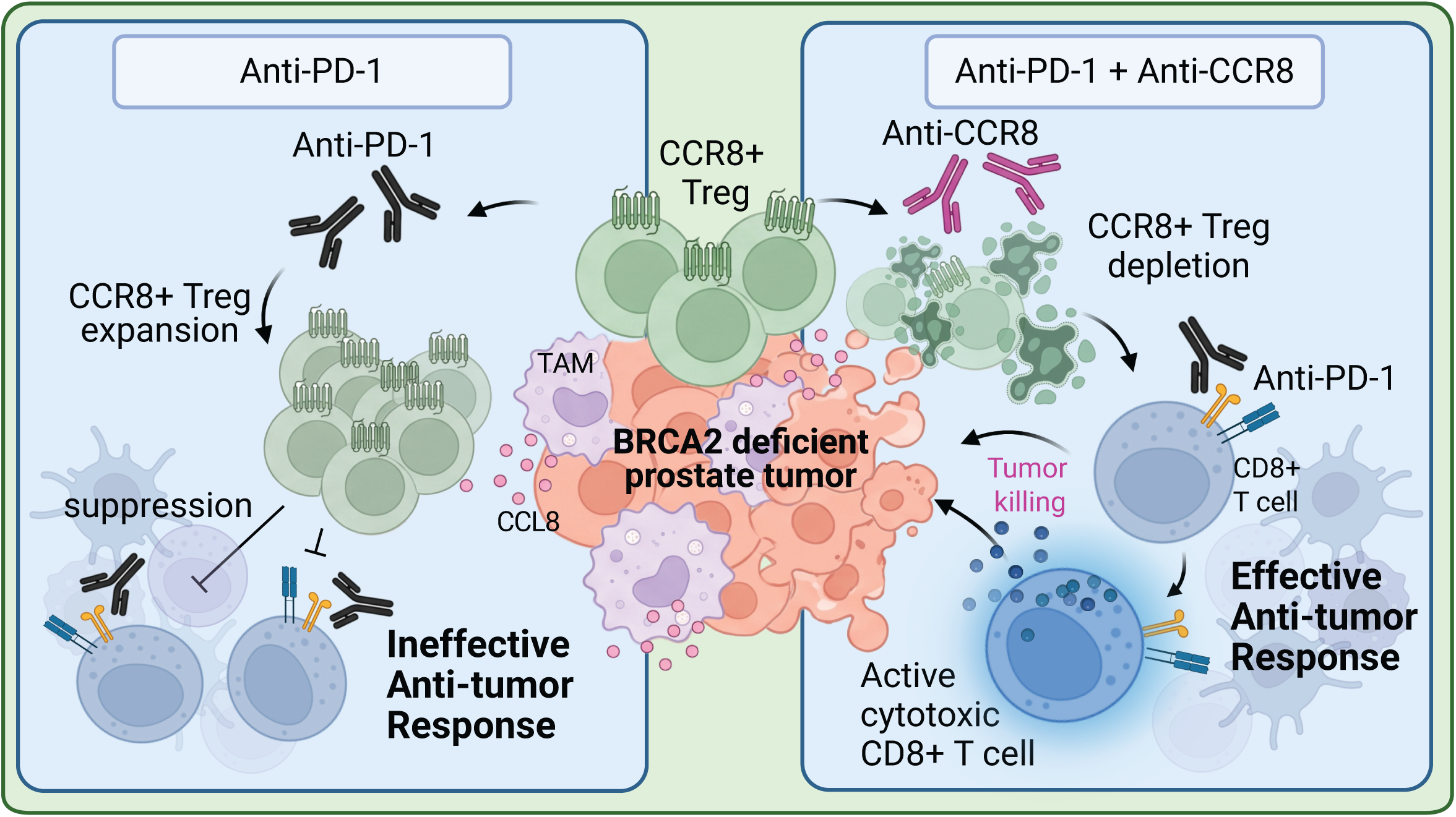
CCR8⁺ regulatory T cells limit checkpoint blockade response in BRCA2-deficient prostate cancer. The proposed model of CCR8⁺ Treg mediated resistance to checkpoint blockade. In *Brca2*-deficient prostate tumors, chronic interferon may promote distinct TAM programming including increased expression of the CCR8 ligand, CCL8, and recruitment of immunosuppressive CCR8⁺ Tregs. Left: anti-PD-1 alone results in expansion of immunosuppressive CCR8⁺ Tregs and the anti-tumor response is ineffective. Right: anti-CCR8 monoclonal antibody depletes CCR8⁺ Tregs, and in combination with anti-PD-1 promotes an effective anti-tumor response with preserved granzyme B⁺ cytotoxic CD8⁺ T cells. CCR8⁺ Treg enrichment and expansion in response to anti-PD-1 is a potential mechanism of checkpoint blockade treatment failure in *Brca2*-deficient prostate cancer. Created in BioRender. Lab, A. (2027) https://BioRender.com/utkfchb.

## Acknowledgements

We thank the UCSF core facilities including Helen Diller Family Comprehensive Cancer Center (HDFCCC) Laboratory for Cell Analysis and Preclinical Core both supported by the NCI/NIH grant P30CA082103, and the UCSF Parnassus Flow CoLab (RRID:SCR_018206). We thank Veronica Steri for her preclinical study support. We thank the UCSF CAT where sequencing was performed and is supported by UCSF PBBR, RRP IMIA, and NIH 1S10OD028511-01 grants. Some results shown here were in part based upon data generated by TCGA Research Network.

## Author Contributions

K.A.R. and A.A. conceived, designed, and supervised the study. K.A.R., I.Q.S., R.P., and V.R. performed experiments. K.A.R., X.Z., E.Y.C., N.Z., and H.S. performed bioinformatics analysis. H.N.V, F.W.H., and M.E.D. contributed to study design and interpretation of findings. C.T. and Y.L. performed Tempus AI cohort data management and curation. K.A.R., A.A., and I.Q.S. wrote the manuscript. All authors reviewed and edited the manuscript and approved the submitted version.

## Competing Interests

A.A. is a co-founder of Azkarra Therapeutics, Ovibio Corporation, Tango Therapeutics and Tiller Tx; a member of the board of Cambridge Science Corporation, Cytomx and Ovibio; a member of the scientific advisory board of Ambagon, Bluestar/Clearnote Health, Circle, Deciphera, Genvivo, GLAdiator, Interdict Bio Inc., Earli, ORIC, Phoenix Molecular Designs, Trial Library, Yingli/280Bio and Z prime; a consultant for Catalio, Longitude, Novartis, ProLynx and YK bioventures; and holds patents on the use of PARP inhibitors held jointly with AstraZeneca from which he has benefited financially (and may do so in the future). C.T. and Y.L. are employees of Tempus AI, Inc. and hold equity in Tempus AI, Inc. The remaining authors declare no competing interests.

## Funding

This research was supported by the UCSF Prostate SPORE Developmental Research Program (P50CA275741), BRCA Research & Cure Alliance (CureBRCA) Grant, and UCSF Benioff Initiative for Prostate Cancer Research. Individual investigators were supported as follows: K.A.R. was supported by NIH T32 Training Grant: Molecular and Cellular Mechanisms in Cancer (T32CA108462), Prostate Cancer Foundation Young Investigator Award, Conquer Cancer ASCO Foundation Young Investigator Award, and UCSF Prostate Cancer Program Pilot award. R.P. was supported by UCSF Rheumatology T32 (5T32AR079068) and the Conquer Cancer ASCO Foundation Young Investigator Award. X.Z. was supported by a Prostate Cancer Foundation Young Investigator award and DoD Physician Research Award (HT9425-24-1-0506). E.Y.C. was supported by the Conquer Cancer ASCO Foundation Young Investigator Award. V.R. was supported by the Fondazione AIRC Fellowship for Abroad Predoc. H.N.V. was supported by a Francis Collins Scholar Award. The remaining authors declare no relevant funding.

## Data Availability

All sequencing data generated for this publication have been deposited in the NCBI Gene Expression Omnibus (GEO) under accession number (pending). Tempus AI human tumor transcriptomic and genomic data were obtained under a data use agreement and are not publicly available; however, requests for access should be directed to Tempus AI. The Cancer Genome Atlas data is publicly available.

## Code Availability

Custom code for mouse sequencing analyses is available at Zenodo under DOI (pending). The analysis was performed in R 4.5.1. Tempus AI cohort analysis was performed within the Tempus Lens environment and can be requested from the corresponding authors and Tempus AI, Inc.

## Ethics

All animal experiments were conducted in accordance with protocols AN206976-00 and AN200569-00, approved by the University of California, San Francisco Institutional Animal Care and Use Committee (IACUC). For the Tempus AI cohort, this study was conducted in accordance with the Declaration of Helsinki, the Belmont Report, and the U.S. Common Rule. In compliance with 45 CFR 46.104(d), the study used retrospective, de-identified clinical data and is considered exempt from institutional review board review, with patient consent not required.

## Methods

### Cell Culture

MyC-CaP cell lines were purchased from ATCC and grown in DMEM (Gibco, 10569044) supplemented with 10% FBS (Corning, MT35010CV) and 1% Penicillin-Streptomycin (Gibco, 15140163) in a humidified incubator at 37 °C and 5% CO_2_. Cells were regularly tested for *Mycoplasma* contamination (Sigma-Aldrich, MP0035-1KT).

### CRISPR/Cas9 Gene Targeting

CRISPR/Cas9-mediated genome editing was performed via electroporation of Cas9/gRNA ribonucleoprotein (RNP) complexes. sgRNA was prepared by combining crRNA and universal tracrRNA, and gRNA duplex was combined with Cas9. RNP complexes were introduced into MyC-CaP cells using the Lonza 4D-Nucleofector system (program EN-120). Post-nucleofection single cells were plated into 96-well plates, expanded, and screened by Sanger sequencing of genomic DNA. Sanger sequencing primers are listed in Supplementary Table 2.

### Cell Survival Assay

Cells were seeded in 96-well plates in triplicate and treated the following day with PARP inhibitors olaparib, talazoparib, rucaparib, niraparib, SN-38, and cisplatin. For cisplatin, the treatment was for 1 h and then washed off and replaced with fresh media. When the untreated wells reached near confluence (∼6-7 days after plating) cells were imaged using the Incucyte Zoom system and relative confluence was calculated.

### RT-qPCR

Total RNA was harvested from mouse tumors or cultured cells using the RNeasy kit (Qiagen, 74106) and quantified on a Nanodrop spectrophotometer system. cDNA was reversed-transcribed from 0.5 µg of total RNA using the PrimeScript RT Master Mix (Takara Bio, RR036A). The iTaq Universal SYBR Green Supermix (BioRad, 1725121) system was used as per manufacturer’s protocol to determine relative transcript levels normalized to GAPDH expression levels. Primer sequences are listed in Supplementary Table 2.

### Whole exome sequencing

Genomic DNA was extracted using the Qiagen DNeasy Blood & Tissue kit. Exome capture and sequencing were performed by MedGenome Inc. (Foster City, CA) using the Agilent SureSelectXT Mouse All Exon kit with paired-end sequencing (2 × 100 bp) on an Illumina NovaSeq 6000/X Plus. Reads were aligned to the mouse reference genome (mm10/GRCm38) using BWA-MEM (v0.7.17-r1188) in Sentieon (v202308.02) and variants called with Sentieon Haplotyper. Indels at the *Brca2* locus were visualized in IGV.

### RNA sequencing

Tumor tissue (10-50 mg) from mice was used fresh or preserved in RNAlater (Invitrogen, AM7020) at -20°C. Tissue was minced in Buffer RLT (RNeasy kit; Qiagen, 74106) with 1% 2-Mercaptoethanol (Sigma-Aldrich, M3148-25ML) and homogenized with Tissue-Tearor (BioSpec Products, Motor, EUR-TT; 9853G-04). Lysates were processed in a QIAshredder column (Qiagen, 79656) to ensure uniform homogenization and RNA was extracted per Qiagen RNeasy kit. Library preparation and sequencing were performed by MedGenome Inc. RNA libraries were prepared using the Illumina Stranded mRNA Prep kit. Paired-end sequencing (2 × 150 bp) was performed on an Illumina NovaSeq platform to a target depth of 20 million paired-end reads per sample. Reads were depleted of contaminating sequences with Bowtie2 (v2.5.3) and aligned to the mouse reference genome (mm10/GRCm38) using STAR (v2.7.11b) and gene-level counts were quantified using HTSeq-count (v2.0.5) against Ensembl gene annotation (release 92).

Raw counts were processed with DESeq2 (v1.48.2).^72^ The DESeq2 model was fit to all samples across genotype comparisons or treatment arms (∼ group) per experiment so that size factors and gene-wise dispersions were estimated jointly. Pairwise contrasts between treatment arms were then extracted. Genes with fewer than 10 counts in at least N samples (N = smallest group size) were excluded prior to fitting. Log2 fold changes shown in the volcano plot were shrunk with ashr (v2.2-63).^73^ For heatmaps, variance-stabilized expression was averaged per treatment arm and row-Z-scored across arms and rendered with ComplexHeatmap (v2.24.1).^74^ Mouse immune-cell gene signatures were assembled from established tumor immunology references and canonical markers.^75,76^ Hallmark pathway enrichment^77^ was performed with the fgsea R package (v1.34.2) on Wald-statistic-ranked genes per contrast, using the mouse MSigDB Hallmark collection accessed via msigdbr (v26.1.0).^78^

### Single-cell RNA sequencing

Syngeneic MyC-CaP tumors, *Brca2^fs^* and *Brca2^WT^*, were dissociated and single-cell libraries prepared using the Fluent BioSciences PIPseq platform. PIPseeker (v3.3.0, chemistry V; Fluent BioSciences GRCm39-2022.04 reference) was used to generate per-sample count matrices. Analysis was performed with Seurat (v5.5.0)^79^ with retention of cells with 500-7,500 detected genes, 1,000-60,000 UMIs, and <15% mitochondrial reads. Doublets were removed with scDblFinder (v1.22.0).^80^ Samples were SCTransform-normalized,^81^ integrated by RPCA, and a clustering resolution of 0.6 was used. Cell types (clusters) were annotated using SingleR (v2.10.0; MouseRNAseqData and ImmGenData references),^82^ followed by manual curation using canonical markers. Differentially expressed genes were computed using FindAllMarkers function in Seurat. Myeloid cells were re-clustered independently to assess the macrophage compartment at resolution 0.5. Differential abundance was assessed using propeller (speckle v1.8.0),^83^ with miloR (v2.4.1)^84^ as a cluster-free sensitivity analysis. Macrophage subcluster abundance was tested using propeller and a negative-binomial GLM; with propeller and GLM in agreement for direction of all comparisons. Pseudobulk profiles were generated by summing raw RNA counts across cells within each sample and cell type. Differential expression between genotypes was performed on sample-level pseudobulk using DESeq2 and Wald-statistic–ranked genes were tested by Hallmark gene-set enrichment (fgsea). Transcription-factor activity was inferred with decoupleR (v2.14.0; univariate linear model)^85^ on the macrophage-compartment pseudobulk DESeq2 Wald statistic using CollecTRI mouse regulons.^86^ Full analysis parameters and code are available at Zenodo DOI 10.5281/zenodo.24024983.

### Animal experiments and preclinical studies

Male FVB/N mice were obtained from Jackson Laboratory at 5–6 weeks of age and housed under standard conditions with food and water available *ad libitum*. Subcutaneous tumors were not permitted to exceed 2,000 mm³ or 2 cm in any dimension. Mice were euthanized by CO₂ asphyxiation. 8.0 x 10^6^ MyC-CaP cells at 1:1 in DMEM and Matrigel (Corning, CB-40234C) were injected subcutaneously in the flank. Mice were weighed once a week, and tumor volumes were measured twice a week by calipers and tumor volume calculated as (length × width²)/2.

Mice were randomized to treatment cohorts and treatments were started when tumors reached ∼100-200 mm^3^ in size. Olaparib was formulated at 2.5 mg/mL in 10% 2-hydroxy-propyl-β-cyclodextrin/PBS with 5% DMSO and administered intraperitoneally at a dose of 75 mg/kg daily. Anti-CCR8 monoclonal antibody (Bioxcell, BE0464) was diluted in PBS and administered intraperitoneally for a dose of 10 mg/kg, 3 times a week (MWF). Anti-PD-1 monoclonal antibody (Bioxcell, BE0146) diluted in PBS and administered intraperitoneally for a dose of 10 mg/kg, 3 times a week (MWF). Talazoparib was formulated at 0.033 mg/mL in 10% DMAc/6% Solutol HS-15/84% PBS with 0.9% DMSO and administered orally by gavage at 0.11 mg/kg or 0.33 mg/kg once daily, 5 days a week (M-F).

### Tumor dissociation and spectral flow cytometry

Tumors from mice were harvested fresh, minced on ice, and digested with 6 mg/mL Collagenase IV. Tumor suspensions were digested at 37°C for 1 hour with agitation every 10 min. Suspensions were filtered through a 40 µm cell strainer. Red blood cells were lysed with 1× RBC Lysis Buffer for 5 minutes. Processed tumor cells were either cryopreserved in FBS with 10% DMSO or stained immediately. Cells were counted and washed with FACS buffer (PBS + Penicillin Streptomycin L-glutamine + 2 mM EDTA + 2% heat-inactivated FCS) twice in a 96-well round-bottom plate. Cells were subsequently stained with surface marker antibodies (Supplementary Table 1) and LIVE/DEAD Fixable NIR stain in FACS buffer for 20 minutes on ice in the dark. Cells were washed twice with FACS buffer and fixed with FoxP3 Fixation-Permeabilization Buffer (Invitrogen 00-5523-00) for 30 minutes at room temperature in the dark. Cells were washed twice with Permeabilization Wash Buffer (Invitrogen 00-5523-00) and stained with intracellular antibodies (Supplementary Table 1) in Permeabilization Wash Buffer for 45 minutes at room temperature in the dark. Cells were then washed twice in Permeabilization Wash Buffer and resuspended in FACS buffer and 4% PFA at a 1:1 ratio for flow cytometry analysis. After staining, cells were analyzed on an Aurora (Cytek) spectral flow cytometer (full spectrum). Single stains were run at the time of each experiment and unmixing was performed via SpectroFlo (v3.3.0) software prior to running samples. Data analysis was performed using FlowJo (v10.10.1) software.

### Western blot analysis

Cells were lysed in Laemmli buffer made up of (4% SDS (Sigma-Aldrich, 05030-500ML-F), 20% glycerol (Sigma-Aldrich, G5516-1L), 120 mM Tris-HCl pH 6.8 (Kd Medical Inc, 50-103-1384)) and sonicated for 10 seconds on ice. Samples were boiled for 7 min, and protein content was quantified by Pierce BCA Protein Assay (ThermoFisher Scientific, 23227). Equal amounts of protein were separated on 4%-12% Bolt Bis-Tris gels (Invitrogen, NW04120BOX) using the Invitrogen system and transferred onto PVDF membranes using the BioRad TransBlot Turbo Transfer System (BioRad, 1704272). Membranes were blocked in 5% milk (Apex Bioresearch Products, 20-241) in TBST. Primary antibodies were incubated at 4°C overnight, secondary antibodies were incubated at room temperature for 1 hour. Antibodies for immunoblotting are listed in Supplementary Table 1.

### Immunofluorescence

Cells were seeded on coverslips overnight. For drug treatments, cells were treated with 2 µM olaparib or 2 nM talazoparib and harvested at indicated timepoint. To induce RAD51 foci, cells were treated with 3 µM MMC for 1 hour and were harvested at indicated timepoint. Cells were washed twice with PBS, fixed with 4% paraformaldehyde, and permeabilized with 0.5% Triton in PBS for 10 minutes. Coverslips were blocked in 5% FBS in PBS and incubated with primary antibodies in blocking buffer for 2 h at room temperature or overnight at 4°C. Cells were incubated with secondary antibodies (Alexa fluor, 1:2500). Coverslips were mounted onto slides with DAPI Fluoromount-G (SouthernBiotech, OB010020). RAD51 foci were imaged using a 63× objective with Zeiss AxioImager M1 and all other slides were imaged using a 40× objective on a Keyence BZ-X800. Antibodies for IF are listed in Supplementary Table 1.

### Tempus AI cohort analysis

The Tempus Lens Platform (Tempus AI, Inc., Chicago, IL) was used to query the multimodal de-identified database and establish a cohort of 1198 patients with prostate cancer. We retrospectively ascertained patients whose prostate tumor harbor pathogenic or likely pathogenic DNA alterations in *BRCA2* (n=271) and patients whose prostate tumor does not harbor any pathogenic or likely pathogenic DNA alteration in *BRCA2* or other HRR genes (*ATM, ATR, ATRX, BAP1, BARD1, BLM, BRCA1, BRIP1, CDK12, CHEK1, CHEK2, FANCA, FANCL, HDAC2, MRE11, NBN, PALB2, RAD50, RAD51B, RAD51C, RAD51D,* and *RAD54L;* n=417). We analyzed the genomic and transcriptomic profiles using *tempusverse* and custom code in R. The Wilcoxon rank-sum test and the Hodges-Lehmann estimator were used to compare *CCR8* and other immune-readout expressions between *BRCA2*-mutated cases and control groups. Mean-z scoring and gene set variation analysis (GSVA; v2.2.1) were used to compute composite checkpoint, IFN, T-cell-inflamed, and curated pathway module scores.^87^ Immune cell-type deconvolution was performed with BayesPrism using a prostate single-cell RNA-seq reference comprising tumor and benign tissue.^88,89^ The Benjamini-Hochberg procedure was used to control the false discovery rate within each exploratory readout family.

### TCGA cohort analysis

TCGA prostate adenocarcinoma (TCGA-PRAD), ovarian cancer (TCGA-OV), and breast cancer (TCGA-BRCA) datasets were downloaded from cBioPortal.^90,91^ For each cohort, samples were stratified by *BRCA2* status (Fig. 6J, TCGA-PRAD) or by *BRCA1* and *BRCA2* status (Fig. 6K, pooled cohorts) using the corresponding somatic mutation annotation .maf files. mRNA expression (RSEM, batch normalized, log2(value+1)) of *CCL18*, and *CCR8* was extracted and compared between the two groups in each cohort. Statistical significance was computed using a two-sided Wilcoxon rank-sum test.

### Statistics

The statistical analyses of the tumor growth and survival, spectral flow cytometry, and immunofluorescence quantification were performed using GraphPad Prism 10. Tests used for each experiment are stated in the figure legend.

### Use of Artificial Intelligence

Artificial intelligence (Claude, Anthropic) was used to assist in writing and editing analysis code. All code was reviewed and executed by authors.

## References

1. Robinson, D., Van Allen, E.M., Wu, Y.M., Schultz, N., Lonigro, R.J., Mosquera, J.M., Montgomery, B., Taplin, M.E., Pritchard, C.C., Attard, G., et al. (2015). Integrative clinical genomics of advanced prostate cancer. Cell 161, 1215–1228. 10.1016/j.cell.2015.05.001.

2. Chung, J.H., Dewal, N., Sokol, E., Mathew, P., Whitehead, R., Millis, S.Z., Frampton, G.M., Bratslavsky, G., Pal, S.K., Lee, R.J., et al. (2019). Prospective Comprehensive Genomic Profiling of Primary and Metastatic Prostate Tumors. JCO Precis Oncol 3. 10.1200/PO.18.00283.

3. Abida, W., Armenia, J., Gopalan, A., Brennan, R., Walsh, M., Barron, D., Danila, D., Rathkopf, D., Morris, M., Slovin, S., et al. (2017). Prospective Genomic Profiling of Prostate Cancer Across Disease States Reveals Germline and Somatic Alterations That May Affect Clinical Decision Making. JCO Precis Oncol 2017. 10.1200/PO.17.00029.

4. Pritchard, C.C., Mateo, J., Walsh, M.F., De Sarkar, N., Abida, W., Beltran, H., Garofalo, A., Gulati, R., Carreira, S., Eeles, R., et al. (2016). Inherited DNA-Repair Gene Mutations in Men with Metastatic Prostate Cancer. N Engl J Med 375, 443–453. 10.1056/NEJMoa1603144.

5. Dhawan, M., Ryan, C.J., and Ashworth, A. (2016). DNA Repair Deficiency Is Common in Advanced Prostate Cancer: New Therapeutic Opportunities. Oncologist 21, 940–945. 10.1634/theoncologist.2016-0135.

6. Castro, E., Goh, C., Leongamornlert, D., Saunders, E., Tymrakiewicz, M., Dadaev, T., Govindasami, K., Guy, M., Ellis, S., Frost, D., et al. (2015). Effect of BRCA Mutations on Metastatic Relapse and Cause-specific Survival After Radical Treatment for Localised Prostate Cancer. Eur Urol 68, 186–193. 10.1016/j.eururo.2014.10.022.

7. Castro, E., Goh, C., Olmos, D., Saunders, E., Leongamornlert, D., Tymrakiewicz, M., Mahmud, N., Dadaev, T., Govindasami, K., Guy, M., et al. (2013). Germline BRCA mutations are associated with higher risk of nodal involvement, distant metastasis, and poor survival outcomes in prostate cancer. J Clin Oncol 31, 1748–1757. 10.1200/JCO.2012.43.1882.

8. Darst, B.F., Saunders, E., Dadaev, T., Sheng, X., Wan, P., Pooler, L., Xia, L.Y., Chanock, S., Berndt, S.I., Wang, Y., et al. (2023). Germline Sequencing Analysis to Inform Clinical Gene Panel Testing for Aggressive Prostate Cancer. JAMA Oncol 9, 1514–1524. 10.1001/jamaoncol.2023.3482.

9. Gallagher, D.J., Gaudet, M.M., Pal, P., Kirchhoff, T., Balistreri, L., Vora, K., Bhatia, J., Stadler, Z., Fine, S.W., Reuter, V., et al. (2010). Germline BRCA mutations denote a clinicopathologic subset of prostate cancer. Clin Cancer Res 16, 2115–2121. 10.1158/1078-0432.CCR-09-2871.

10. Moynahan, M.E., Pierce, A.J., and Jasin, M. (2001). BRCA2 is required for homology-directed repair of chromosomal breaks. Mol Cell 7, 263–272. 10.1016/s1097-2765(01)00174-5.

11. Scully, R., Panday, A., Elango, R., and Willis, N.A. (2019). DNA double-strand break repair-pathway choice in somatic mammalian cells. Nat Rev Mol Cell Biol 20, 698–714. 10.1038/s41580-019-0152-0.

12. Schlacher, K., Christ, N., Siaud, N., Egashira, A., Wu, H., and Jasin, M. (2011). Double-strand break repair-independent role for BRCA2 in blocking stalled replication fork degradation by MRE11. Cell 145, 529–542. 10.1016/j.cell.2011.03.041.

13. Lord, C.J., Tutt, A.N.J., and Ashworth, A. (2026). Two decades of PARP inhibitor synthetic lethality in cancer. Nature 653, 41–51. 10.1038/s41586-026-10404-y.

14. de Bono, J., Mateo, J., Fizazi, K., Saad, F., Shore, N., Sandhu, S., Chi, K.N., Sartor, O., Agarwal, N., Olmos, D., et al. (2020). Olaparib for Metastatic Castration-Resistant Prostate Cancer. N Engl J Med 382, 2091–2102. 10.1056/NEJMoa1911440.

15. Abida, W., Patnaik, A., Campbell, D., Shapiro, J., Bryce, A.H., McDermott, R., Sautois, B., Vogelzang, N.J., Bambury, R.M., Voog, E., et al. (2020). Rucaparib in Men With Metastatic Castration-Resistant Prostate Cancer Harboring a BRCA1 or BRCA2 Gene Alteration. J Clin Oncol 38, 3763–3772. 10.1200/JCO.20.01035.

16. Attard, G., Agarwal, N., Graff, J.N., Sandhu, S., Efstathiou, E., Ozguroglu, M., Pereira de Santana Gomes, A.J., Vianna, K., Luo, H., Gotto, G.T., et al. (2025). Niraparib and abiraterone acetate plus prednisone for HRR-deficient metastatic castration-sensitive prostate cancer: a randomized phase 3 trial. Nat Med 31, 4109–4118. 10.1038/s41591-025-03961-8.

17. Agarwal, N., Matsubara, N., Azad, A.A., Saad, F., Mateo, J., Jiang, S., Ye, D., Voog, E., Shore, N.D., Cil, T., et al. (2026). PARP and Androgen-Signaling Inhibition plus ADT in Metastatic Prostate Cancer. N Engl J Med 395, 427–439. 10.1056/NEJMoa2604126.

18. Abida, W., Cheng, M.L., Armenia, J., Middha, S., Autio, K.A., Vargas, H.A., Rathkopf, D., Morris, M.J., Danila, D.C., Slovin, S.F., et al. (2019). Analysis of the Prevalence of Microsatellite Instability in Prostate Cancer and Response to Immune Checkpoint Blockade. JAMA Oncol 5, 471–478. 10.1001/jamaoncol.2018.5801.

19. Antonarakis, E.S., Piulats, J.M., Gross-Goupil, M., Goh, J., Ojamaa, K., Hoimes, C.J., Vaishampayan, U., Berger, R., Sezer, A., Alanko, T., et al. (2020). Pembrolizumab for Treatment-Refractory Metastatic Castration-Resistant Prostate Cancer: Multicohort, Open-Label Phase II KEYNOTE-199 Study. J Clin Oncol 38, 395–405. 10.1200/JCO.19.01638.

20. Shenderov, E., Boudadi, K., Fu, W., Wang, H., Sullivan, R., Jordan, A., Dowling, D., Harb, R., Schonhoft, J., Jendrisak, A., et al. (2021). Nivolumab plus ipilimumab, with or without enzalutamide, in AR-V7-expressing metastatic castration-resistant prostate cancer: A phase-2 nonrandomized clinical trial. Prostate 81, 326–338. 10.1002/pros.24110.

21. Slovin, S.F., Higano, C.S., Hamid, O., Tejwani, S., Harzstark, A., Alumkal, J.J., Scher, H.I., Chin, K., Gagnier, P., McHenry, M.B., and Beer, T.M. (2013). Ipilimumab alone or in combination with radiotherapy in metastatic castration-resistant prostate cancer: results from an open-label, multicenter phase I/II study. Ann Oncol 24, 1813–1821. 10.1093/annonc/mdt107.

22. Lyu, A., Fan, Z., Clark, M., Lea, A., Luong, D., Setayesh, A., Starzinski, A., Wolters, R., Arias-Badia, M., Allaire, K., et al. (2025). Evolution of myeloid-mediated immunotherapy resistance in prostate cancer. Nature 637, 1207–1217. 10.1038/s41586-024-08290-3.

23. Calcinotto, A., Spataro, C., Zagato, E., Di Mitri, D., Gil, V., Crespo, M., De Bernardis, G., Losa, M., Mirenda, M., Pasquini, E., et al. (2018). IL-23 secreted by myeloid cells drives castration-resistant prostate cancer. Nature 559, 363–369. 10.1038/s41586-018-0266-0.

24. Lopez-Bujanda, Z.A., Haffner, M.C., Chaimowitz, M.G., Chowdhury, N., Venturini, N.J., Patel, R.A., Obradovic, A., Hansen, C.S., Jackow, J., Maynard, J.P., et al. (2021). Castration-mediated IL-8 promotes myeloid infiltration and prostate cancer progression. Nat Cancer 2, 803–818. 10.1038/s43018-021-00227-3.

25. Escamilla, J., Schokrpur, S., Liu, C., Priceman, S.J., Moughon, D., Jiang, Z., Pouliot, F., Magyar, C., Sung, J.L., Xu, J., et al. (2015). CSF1 receptor targeting in prostate cancer reverses macrophage-mediated resistance to androgen blockade therapy. Cancer Res 75, 950–962. 10.1158/0008-5472.CAN-14-0992.

26. Kfoury, Y., Baryawno, N., Severe, N., Mei, S., Gustafsson, K., Hirz, T., Brouse, T., Scadden, E.W., Igolkina, A.A., Kokkaliaris, K., et al. (2021). Human prostate cancer bone metastases have an actionable immunosuppressive microenvironment. Cancer Cell 39, 1464–1478 e1468. 10.1016/j.ccell.2021.09.005.

27. Obradovic, A.Z., Dallos, M.C., Zahurak, M.L., Partin, A.W., Schaeffer, E.M., Ross, A.E., Allaf, M.E., Nirschl, T.R., Liu, D., Chapman, C.G., et al. (2020). T-Cell Infiltration and Adaptive Treg Resistance in Response to Androgen Deprivation With or Without Vaccination in Localized Prostate Cancer. Clin Cancer Res 26, 3182–3192. 10.1158/1078-0432.CCR-19-3372.

28. Erlandsson, A., Carlsson, J., Lundholm, M., Falt, A., Andersson, S.O., Andren, O., and Davidsson, S. (2019). M2 macrophages and regulatory T cells in lethal prostate cancer. Prostate 79, 363–369. 10.1002/pros.23742.

29. Sfanos, K.S., Bruno, T.C., Maris, C.H., Xu, L., Thoburn, C.J., DeMarzo, A.M., Meeker, A.K., Isaacs, W.B., and Drake, C.G. (2008). Phenotypic analysis of prostate-infiltrating lymphocytes reveals TH17 and Treg skewing. Clin Cancer Res 14, 3254–3261. 10.1158/1078-0432.CCR-07-5164.

30. Davidsson, S., Ohlson, A.L., Andersson, S.O., Fall, K., Meisner, A., Fiorentino, M., Andren, O., and Rider, J.R. (2013). CD4 helper T cells, CD8 cytotoxic T cells, and FOXP3(+) regulatory T cells with respect to lethal prostate cancer. Mod Pathol 26, 448–455. 10.1038/modpathol.2012.164.

31. Stultz, J., and Fong, L. (2021). How to turn up the heat on the cold immune microenvironment of metastatic prostate cancer. Prostate Cancer Prostatic Dis 24, 697–717. 10.1038/s41391-021-00340-5.

32. Bezzi, M., Seitzer, N., Ishikawa, T., Reschke, M., Chen, M., Wang, G., Mitchell, C., Ng, C., Katon, J., Lunardi, A., et al. (2018). Diverse genetic-driven immune landscapes dictate tumor progression through distinct mechanisms. Nat Med 24, 165–175. 10.1038/nm.4463.

33. Wu, Y.M., Cieslik, M., Lonigro, R.J., Vats, P., Reimers, M.A., Cao, X., Ning, Y., Wang, L., Kunju, L.P., de Sarkar, N., et al. (2018). Inactivation of CDK12 Delineates a Distinct Immunogenic Class of Advanced Prostate Cancer. Cell 173, 1770–1782 e1714. 10.1016/j.cell.2018.04.034.

34. Mackenzie, K.J., Carroll, P., Martin, C.A., Murina, O., Fluteau, A., Simpson, D.J., Olova, N., Sutcliffe, H., Rainger, J.K., Leitch, A., et al. (2017). cGAS surveillance of micronuclei links genome instability to innate immunity. Nature 548, 461–465. 10.1038/nature23449.

35. Harding, S.M., Benci, J.L., Irianto, J., Discher, D.E., Minn, A.J., and Greenberg, R.A. (2017). Mitotic progression following DNA damage enables pattern recognition within micronuclei. Nature 548, 466–470. 10.1038/nature23470.

36. van Vugt, M., and Parkes, E.E. (2022). When breaks get hot: inflammatory signaling in BRCA1/2-mutant cancers. Trends Cancer 8, 174–189. 10.1016/j.trecan.2021.12.003.

37. Sun, L., Wu, J., Du, F., Chen, X., and Chen, Z.J. (2013). Cyclic GMP-AMP synthase is a cytosolic DNA sensor that activates the type I interferon pathway. Science 339, 786–791. 10.1126/science.1232458.

38. Ishikawa, H., Ma, Z., and Barber, G.N. (2009). STING regulates intracellular DNA-mediated, type I interferon-dependent innate immunity. Nature 461, 788–792. 10.1038/nature08476.

39. Bakhoum, S.F., Ngo, B., Laughney, A.M., Cavallo, J.A., Murphy, C.J., Ly, P., Shah, P., Sriram, R.K., Watkins, T.B.K., Taunk, N.K., et al. (2018). Chromosomal instability drives metastasis through a cytosolic DNA response. Nature 553, 467–472. 10.1038/nature25432.

40. Kwon, J., and Bakhoum, S.F. (2020). The Cytosolic DNA-Sensing cGAS-STING Pathway in Cancer. Cancer Discov 10, 26–39. 10.1158/2159-8290.CD-19-0761.

41. Ahn, J., Xia, T., Konno, H., Konno, K., Ruiz, P., and Barber, G.N. (2014). Inflammation-driven carcinogenesis is mediated through STING. Nat Commun 5, 5166. 10.1038/ncomms6166.

42. Li, Q., Song, Q., Ma, L., Kang, K., Zhu, X., Li, Y., Lin, X., Lu, X., Zeng, Z., Li, G., et al. (2026). The cGAS-STING pathway in cancer: friend or foe. Cell Death Dis 17. 10.1038/s41419-026-08607-2.

43. Lord, C.J., and Ashworth, A. (2016). BRCAness revisited. Nat Rev Cancer 16, 110–120. 10.1038/nrc.2015.21.

44. Reislander, T., Lombardi, E.P., Groelly, F.J., Miar, A., Porru, M., Di Vito, S., Wright, B., Lockstone, H., Biroccio, A., Harris, A., et al. (2019). BRCA2 abrogation triggers innate immune responses potentiated by treatment with PARP inhibitors. Nat Commun 10, 3143. 10.1038/s41467-019-11048-5.

45. Heijink, A.M., Talens, F., Jae, L.T., van Gijn, S.E., Fehrmann, R.S.N., Brummelkamp, T.R., and van Vugt, M. (2019). BRCA2 deficiency instigates cGAS-mediated inflammatory signaling and confers sensitivity to tumor necrosis factor-alpha-mediated cytotoxicity. Nat Commun 10, 100. 10.1038/s41467-018-07927-y.

46. Concannon, K., Morris, B.B., Gay, C.M., and Byers, L.A. (2023). Combining targeted DNA repair inhibition and immune-oncology approaches for enhanced tumor control. Mol Cell 83, 660–680. 10.1016/j.molcel.2022.12.031.

47. Watson, P.A., Ellwood-Yen, K., King, J.C., Wongvipat, J., Lebeau, M.M., and Sawyers, C.L. (2005). Context-dependent hormone-refractory progression revealed through characterization of a novel murine prostate cancer cell line. Cancer Res 65, 11565–11571. 10.1158/0008-5472.CAN-05-3441.

48. Farmer, H., McCabe, N., Lord, C.J., Tutt, A.N., Johnson, D.A., Richardson, T.B., Santarosa, M., Dillon, K.J., Hickson, I., Knights, C., et al. (2005). Targeting the DNA repair defect in BRCA mutant cells as a therapeutic strategy. Nature 434, 917–921. 10.1038/nature03445.

49. Cords, L., Tietscher, S., Anzeneder, T., Langwieder, C., Rees, M., de Souza, N., and Bodenmiller, B. (2023). Cancer-associated fibroblast classification in single-cell and spatial proteomics data. Nat Commun 14, 4294. 10.1038/s41467-023-39762-1.

50. Cumming, J., Maneshi, P., Dongre, M., Alsaed, T., Dehghan-Nayeri, M.J., Ling, A., Pietras, K., Patthey, C., and Ohlund, D. (2025). Dissecting FAP+ Cell Diversity in Pancreatic Cancer Uncovers an Interferon-Response Subtype of Cancer-Associated Fibroblasts with Tumor-Restraining Properties. Cancer Res 85, 2388–2411. 10.1158/0008-5472.CAN-23-3252.

51. Campisi, M., Sundararaman, S.K., Shelton, S.E., Knelson, E.H., Mahadevan, N.R., Yoshida, R., Tani, T., Ivanova, E., Canadas, I., Osaki, T., et al. (2020). Tumor-Derived cGAMP Regulates Activation of the Vasculature. Front Immunol 11, 2090. 10.3389/fimmu.2020.02090.

52. Yang, H., Lee, W.S., Kong, S.J., Kim, C.G., Kim, J.H., Chang, S.K., Kim, S., Kim, G., Chon, H.J., and Kim, C. (2019). STING activation reprograms tumor vasculatures and synergizes with VEGFR2 blockade. J Clin Invest 129, 4350–4364. 10.1172/JCI125413.

53. Islam, S.A., Chang, D.S., Colvin, R.A., Byrne, M.H., McCully, M.L., Moser, B., Lira, S.A., Charo, I.F., and Luster, A.D. (2011). Mouse CCL8, a CCR8 agonist, promotes atopic dermatitis by recruiting IL-5+ T(H)2 cells. Nat Immunol 12, 167–177. 10.1038/ni.1984.

54. Plitas, G., Konopacki, C., Wu, K., Bos, P.D., Morrow, M., Putintseva, E.V., Chudakov, D.M., and Rudensky, A.Y. (2016). Regulatory T Cells Exhibit Distinct Features in Human Breast Cancer. Immunity 45, 1122–1134. 10.1016/j.immuni.2016.10.032.

55. De Simone, M., Arrigoni, A., Rossetti, G., Gruarin, P., Ranzani, V., Politano, C., Bonnal, R.J.P., Provasi, E., Sarnicola, M.L., Panzeri, I., et al. (2016). Transcriptional Landscape of Human Tissue Lymphocytes Unveils Uniqueness of Tumor-Infiltrating T Regulatory Cells. Immunity 45, 1135–1147. 10.1016/j.immuni.2016.10.021.

56. Huang, L., Wang, J., Fang, B., Meric-Bernstam, F., Roth, J.A., and Ha, M.J. (2022). CombPDX: a unified statistical framework for evaluating drug synergism in patient-derived xenografts. Sci Rep 12, 12984. 10.1038/s41598-022-16933-6.

57. Pantelidou, C., Sonzogni, O., De Oliveria Taveira, M., Mehta, A.K., Kothari, A., Wang, D., Visal, T., Li, M.K., Pinto, J., Castrillon, J.A., et al. (2019). PARP Inhibitor Efficacy Depends on CD8(+) T-cell Recruitment via Intratumoral STING Pathway Activation in BRCA-Deficient Models of Triple-Negative Breast Cancer. Cancer Discov 9, 722–737. 10.1158/2159-8290.CD-18-1218.

58. Ding, L., Kim, H.J., Wang, Q., Kearns, M., Jiang, T., Ohlson, C.E., Li, B.B., Xie, S., Liu, J.F., Stover, E.H., et al. (2018). PARP Inhibition Elicits STING-Dependent Antitumor Immunity in Brca1-Deficient Ovarian Cancer. Cell Rep 25, 2972–2980 e2975. 10.1016/j.celrep.2018.11.054.

59. Liang, H., Deng, L., Hou, Y., Meng, X., Huang, X., Rao, E., Zheng, W., Mauceri, H., Mack, M., Xu, M., et al. (2017). Host STING-dependent MDSC mobilization drives extrinsic radiation resistance. Nat Commun 8, 1736. 10.1038/s41467-017-01566-5.

60. Parkes, E.E., Walker, S.M., Taggart, L.E., McCabe, N., Knight, L.A., Wilkinson, R., McCloskey, K.D., Buckley, N.E., Savage, K.I., Salto-Tellez, M., et al. (2017). Activation of STING-Dependent Innate Immune Signaling By S-Phase-Specific DNA Damage in Breast Cancer. J Natl Cancer Inst 109. 10.1093/jnci/djw199.

61. Islam, S.A., Ling, M.F., Leung, J., Shreffler, W.G., and Luster, A.D. (2013). Identification of human CCR8 as a CCL18 receptor. J Exp Med 210, 1889–1898. 10.1084/jem.20130240.

62. Honda, F., Tsuboi, H., Ono, Y., Abe, S., Takahashi, H., Ito, K., Yamada, K., Kawano, M., Kondo, Y., Asano, K., et al. (2021). Pathogenic roles and therapeutic potential of the CCL8-CCR8 axis in a murine model of IgG4-related sialadenitis. Arthritis Res Ther 23, 214. 10.1186/s13075-021-02597-6.

63. Liu, L., Rangan, L., Vanalken, N., Kong, Q., Schlenner, S., De Jonghe, S., Schols, D., and Van Loy, T. (2024). Development of a cellular model to study CCR8 signaling in tumor-infiltrating regulatory T cells. Cancer Immunol Immunother 73, 11. 10.1007/s00262-023-03607-z.

64. Hayashi, Y., Ueyama, A., Funaki, S., Jinushi, K., Higuchi, N., Morihara, H., Hirata, M., Nagira, Y., Saito, T., Kawashima, A., et al. (2024). In situ analysis of CCR8(+) regulatory T cells in lung cancer: suppression of GzmB(+) CD8(+) T cells and prognostic marker implications. BMC Cancer 24, 627. 10.1186/s12885-024-12363-x.

65. Haruna, M., Ueyama, A., Yamamoto, Y., Hirata, M., Goto, K., Yoshida, H., Higuchi, N., Yoshida, T., Kidani, Y., Nakamura, Y., et al. (2022). The impact of CCR8+ regulatory T cells on cytotoxic T cell function in human lung cancer. Sci Rep 12, 5377. 10.1038/s41598-022-09458-5.

66. Jinushi, K., Hayashi, Y., Morishita, K., Saito, T., Kawashima, A., Doki, Y., Ueyama, A., and Wada, H. (2025). The immunosuppressive role of CCR8(+) Tregs in gastric cancer. J Cancer 16, 4390–4399. 10.7150/jca.121363.

67. Kamada, T., Togashi, Y., Tay, C., Ha, D., Sasaki, A., Nakamura, Y., Sato, E., Fukuoka, S., Tada, Y., Tanaka, A., et al. (2019). PD-1(+) regulatory T cells amplified by PD-1 blockade promote hyperprogression of cancer. Proc Natl Acad Sci U S A 116, 9999–10008. 10.1073/pnas.1822001116.

68. van Gulijk, M., van Krimpen, A., Schetters, S., Eterman, M., van Elsas, M., Mankor, J., Klaase, L., de Bruijn, M., van Nimwegen, M., van Tienhoven, T., et al. (2023). PD-L1 checkpoint blockade promotes regulatory T cell activity that underlies therapy resistance. Sci Immunol 8, eabn6173. 10.1126/sciimmunol.abn6173.

69. Geels, S.N., Moshensky, A., Sousa, R.S., Murat, C., Bustos, M.A., Walker, B.L., Singh, R., Harbour, S.N., Gutierrez, G., Hwang, M., et al. (2024). Interruption of the intratumor CD8(+) T cell:Treg crosstalk improves the efficacy of PD-1 immunotherapy. Cancer Cell 42, 1051–1066 e1057. 10.1016/j.ccell.2024.05.013.

70. Wen, Y., Xia, Y., Yang, X., Li, H., and Gao, Q. (2025). CCR8: a promising therapeutic target against tumor-infiltrating regulatory T cells. Trends Immunol 46, 153–165. 10.1016/j.it.2025.01.001.

71. Bockman, A., Zhang, C., Gittins, B., Hung, J., Campbell, T.F., Moreno Ayala, M.A., Saddier Axe, D., Weist, B.M., and DuPage, M. (2026). Intratumoral Regulatory T-cell Ablation Mediates Tumor Control Systemically without Autoimmunity. Cancer Res 86, 4028–4042. 10.1158/0008-5472.CAN-25-3473.

72. Love, M.I., Huber, W., and Anders, S. (2014). Moderated estimation of fold change and dispersion for RNA-seq data with DESeq2. Genome Biol 15, 550. 10.1186/s13059-014-0550-8.

73. Stephens, M. (2017). False discovery rates: a new deal. Biostatistics 18, 275–294. 10.1093/biostatistics/kxw041.

74. Gu, Z., Eils, R., and Schlesner, M. (2016). Complex heatmaps reveal patterns and correlations in multidimensional genomic data. Bioinformatics 32, 2847–2849. 10.1093/bioinformatics/btw313.

75. Bindea, G., Mlecnik, B., Tosolini, M., Kirilovsky, A., Waldner, M., Obenauf, A.C., Angell, H., Fredriksen, T., Lafontaine, L., Berger, A., et al. (2013). Spatiotemporal dynamics of intratumoral immune cells reveal the immune landscape in human cancer. Immunity 39, 782–795. 10.1016/j.immuni.2013.10.003.

76. Charoentong, P., Finotello, F., Angelova, M., Mayer, C., Efremova, M., Rieder, D., Hackl, H., and Trajanoski, Z. (2017). Pan-cancer Immunogenomic Analyses Reveal Genotype-Immunophenotype Relationships and Predictors of Response to Checkpoint Blockade. Cell Rep 18, 248–262. 10.1016/j.celrep.2016.12.019.

77. Subramanian, A., Tamayo, P., Mootha, V.K., Mukherjee, S., Ebert, B.L., Gillette, M.A., Paulovich, A., Pomeroy, S.L., Golub, T.R., Lander, E.S., and Mesirov, J.P. (2005). Gene set enrichment analysis: a knowledge-based approach for interpreting genome-wide expression profiles. Proc Natl Acad Sci U S A 102, 15545–15550. 10.1073/pnas.0506580102.

78. Liberzon, A., Birger, C., Thorvaldsdottir, H., Ghandi, M., Mesirov, J.P., and Tamayo, P. (2015). The Molecular Signatures Database (MSigDB) hallmark gene set collection. Cell Syst 1, 417–425. 10.1016/j.cels.2015.12.004.

79. Hao, Y., Stuart, T., Kowalski, M.H., Choudhary, S., Hoffman, P., Hartman, A., Srivastava, A., Molla, G., Madad, S., Fernandez-Granda, C., and Satija, R. (2024). Dictionary learning for integrative, multimodal and scalable single-cell analysis. Nat Biotechnol 42, 293–304. 10.1038/s41587-023-01767-y.

80. Germain, P.L., Lun, A., Garcia Meixide, C., Macnair, W., and Robinson, M.D. (2021). Doublet identification in single-cell sequencing data using scDblFinder. F1000Res 10, 979. 10.12688/f1000research.73600.2.

81. Hafemeister, C., and Satija, R. (2019). Normalization and variance stabilization of single-cell RNA-seq data using regularized negative binomial regression. Genome Biol 20, 296. 10.1186/s13059-019-1874-1.

82. Aran, D., Looney, A.P., Liu, L., Wu, E., Fong, V., Hsu, A., Chak, S., Naikawadi, R.P., Wolters, P.J., Abate, A.R., et al. (2019). Reference-based analysis of lung single-cell sequencing reveals a transitional profibrotic macrophage. Nat Immunol 20, 163–172. 10.1038/s41590-018-0276-y.

83. Phipson, B., Sim, C.B., Porrello, E.R., Hewitt, A.W., Powell, J., and Oshlack, A. (2022). propeller: testing for differences in cell type proportions in single cell data. Bioinformatics 38, 4720–4726. 10.1093/bioinformatics/btac582.

84. Dann, E., Henderson, N.C., Teichmann, S.A., Morgan, M.D., and Marioni, J.C. (2022). Differential abundance testing on single-cell data using k-nearest neighbor graphs. Nat Biotechnol 40, 245–253. 10.1038/s41587-021-01033-z.

85. Badia, I.M.P., Velez Santiago, J., Braunger, J., Geiss, C., Dimitrov, D., Muller-Dott, S., Taus, P., Dugourd, A., Holland, C.H., Ramirez Flores, R.O., and Saez-Rodriguez, J. (2022). decoupleR: ensemble of computational methods to infer biological activities from omics data. Bioinform Adv 2, vbac016. 10.1093/bioadv/vbac016.

86. Muller-Dott, S., Tsirvouli, E., Vazquez, M., Ramirez Flores, R.O., Badia, I.M.P., Fallegger, R., Turei, D., Laegreid, A., and Saez-Rodriguez, J. (2023). Expanding the coverage of regulons from high-confidence prior knowledge for accurate estimation of transcription factor activities. Nucleic Acids Res 51, 10934–10949. 10.1093/nar/gkad841.

87. Hanzelmann, S., Castelo, R., and Guinney, J. (2013). GSVA: gene set variation analysis for microarray and RNA-seq data. BMC Bioinformatics 14, 7. 10.1186/1471-2105-14-7.

88. Chu, T., Wang, Z., Pe’er, D., and Danko, C.G. (2022). Cell type and gene expression deconvolution with BayesPrism enables Bayesian integrative analysis across bulk and single-cell RNA sequencing in oncology. Nat Cancer 3, 505–517. 10.1038/s43018-022-00356-3.

89. Hirz, T., Mei, S., Sarkar, H., Kfoury, Y., Wu, S., Verhoeven, B.M., Subtelny, A.O., Zlatev, D.V., Wszolek, M.W., Salari, K., et al. (2023). Dissecting the immune suppressive human prostate tumor microenvironment via integrated single-cell and spatial transcriptomic analyses. Nat Commun 14, 663. 10.1038/s41467-023-36325-2.

90. Cerami, E., Gao, J., Dogrusoz, U., Gross, B.E., Sumer, S.O., Aksoy, B.A., Jacobsen, A., Byrne, C.J., Heuer, M.L., Larsson, E., et al. (2012). The cBio cancer genomics portal: an open platform for exploring multidimensional cancer genomics data. Cancer Discov 2, 401–404. 10.1158/2159-8290.CD-12-0095.

91. Gao, J., Aksoy, B.A., Dogrusoz, U., Dresdner, G., Gross, B., Sumer, S.O., Sun, Y., Jacobsen, A., Sinha, R., Larsson, E., et al. (2013). Integrative analysis of complex cancer genomics and clinical profiles using the cBioPortal. Sci Signal 6, pl1. 10.1126/scisignal.2004088.

